# Pervasive contingency and entrenchment in a billion years of Hsp90 evolution

**DOI:** 10.1101/189803

**Authors:** Tyler N. Starr, Julia M. Flynn, Parul Mishra, Daniel N. A. Bolon, Joseph W. Thornton

## Abstract

Interactions among mutations within a protein have the potential to make molecular evolution contingent and irreversible, but the extent to which epistasis actually shaped historical evolutionary trajectories is unclear. We addressed this question by identifying all amino acid substitutions that occurred during the billion-year evolutionary history of the heat shock protein 90 (Hsp90) ATPase domain beginning from a deep eukaryotic ancestor to modern *Saccharomyces cerevisiae* and then precisely measuring their fitness effects when introduced into both extant and reconstructed ancestral Hsp90 proteins. We find a pervasive influence of epistasis: of 98 derived states that evolved during history, most were deleterious at times before they happened, and the vast majority also became subsequently entrenched, with the ancestral state becoming deleterious after its substitution. This epistasis was primarily caused by specific interactions among sites rather than a general permissive or restrictive effect on the protein’s tolerance to mutation. Our results show that epistasis continually opens and closes windows of mutational opportunity over evolutionary timescales, producing histories and biological states that reflect the transient internal constraints imposed by a protein’s fleeting sequence states.

**Significance statement:** When mutations within a protein change each other’s functional effects—a phenomenon called epistasis—the trajectories available to evolution at any moment in time depend on the specific set of changes that previously occurred in the protein. The extent to which epistasis has shaped historical evolutionary trajectories is unknown. Using a high-precision bulk fitness assay and ancestral protein reconstruction, we measured the fitness effects in ancestral and extant sequences of all historical substitutions that occurred during the billion-year trajectory of an essential protein. We found that most historical substitutions were contingent on prior epistatic substitutions and/or entrenched by subsequent changes. These results establish that epistasis caused widespread, consequential shifts in the site-specific fitness constraints that shaped the protein’s historical trajectory.

## Main text

Epistatic interactions can, in principle, affect the sequence changes that accumulate during evolution. A deleterious mutation’s expected fate is to be purged by purifying selection, but it can be fixed if a permissive substitution renders it neutral or beneficial (1-3). Conversely, a neutral mutation – which by definition is initially reversible to the ancestral state without fitness cost – may become entrenched by a subsequent restrictive substitution that renders the ancestral state deleterious (1, 4, 5); reversal of the entrenched mutation would then be unlikely unless the restrictive substitution were itself reversed or another permissive substitution occurred.

The extent to which epistasis-induced contingency and entrenchment actually affected protein sequence evolution remains unclear, however, because there is no consensus on the prevalence, effect size, or mechanisms of epistasis among historical substitutions. Deep mutational scans have revealed frequent epistasis among the many possible mutations within proteins (6-10), but how these interactions affect the substitutions that actually occurred during historical evolution is not known. Historical case studies have shown that particular substitutions were contingent (3, 11-13) or became entrenched during evolution (5), but whether these are examples of a general phenomenon is unknown. Computational approaches suggest pervasive contingency and entrenchment among substitutions (1, 4, 14-18), but some of these analyses rely on models of uncertain adequacy (19-21), and their claims have not been experimentally validated. Swapping sequence states among extant orthologs reveals frequent epistasis among substitutions (22), but this “horizontal” approach, unpolarized with respect to time, leaves unresolved whether permissive or restrictive interactions are at play (23). Some experimental studies have systematically examined epistasis among substitutions in an historical context, but most have measured effects on protein function (2, 22) or stability (20, 24), leaving unexamined the prevalence of epistasis with respect to fitness – the phenotype that directly affects evolutionary fate. Others have focused on fitness but used methods that cannot detect effects of relatively small magnitude, which could be both widespread and consequential for evolutionary processes (2, 25).

We directly evaluated the roles of contingency and entrenchment on historical sequence evolution by precisely quantifying changes over time in the fitness effects of all substitutions that accumulated during the long-term evolution of heat shock protein 90 (Hsp90) from a deep eukaryotic ancestor to *S. cerevisiae*. Hsp90 is an essential molecular chaperone that facilitates folding and regulation of substrate proteins through an ATP-dependent cycle of conformational changes, modulated by co-chaperone proteins. Orthologs from other fungi, animals, and protists can complement Hsp90 deletion in *S. cerevisiae* (26, 27), indicating that the protein’s essential molecular function is conserved over large evolutionary distances. To quantify the context-dependence of historical sequence changes, we used a sensitive deep sequencing-based bulk fitness assay (28) to characterize protein libraries in which each ancestral amino acid is reintroduced into an extant Hsp90 and each derived state is introduced into a reconstructed ancestral Hsp90. We focused our experiments on the N-terminal domain (NTD) of Hsp90, which mediates ATP-dependent conformational changes.

## Results

### The historical trajectory of Hsp90 sequence evolution

We inferred the maximum likelihood phylogeny of Hsp90 protein sequences from 261 species of Amorphea (the clade comprising Fungi, Metazoa, Amoebozoa, and related lineages (29)), rooted using green algae and plants as an outgroup (Fig. 1a, Fig. S1, Datasets S1, S2, S3). We reconstructed ancestral NTD sequences at all nodes along the trajectory from the common ancestor of Amorphea (ancAmoHsp90) to extant *S. cerevisiae* (ScHsp90) and identified substitutions as differences between the most probable reconstructions at successive nodes (Dataset S4).

**Figure 1.**
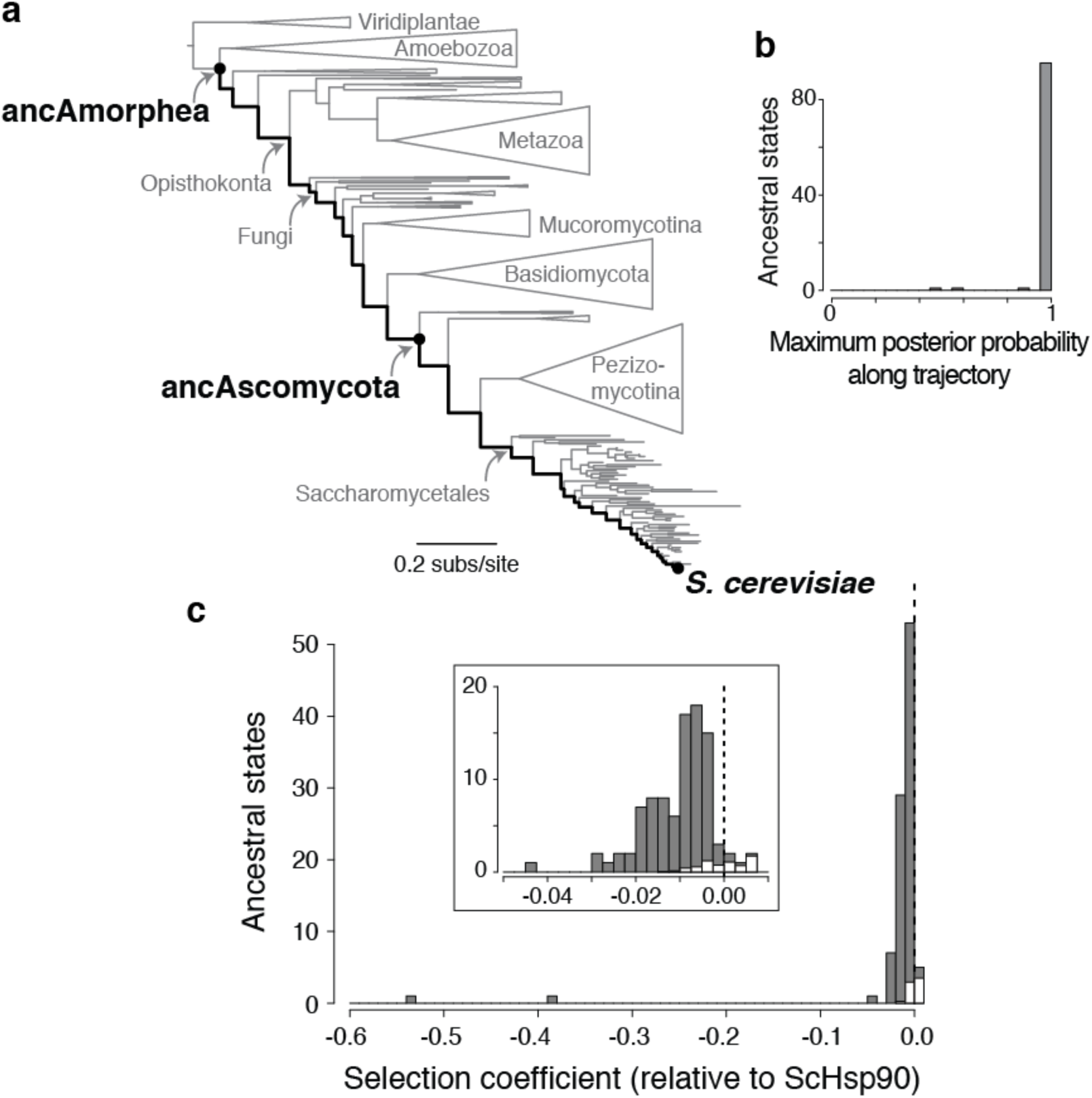
Ancestral states are deleterious in the yeast Hsp90 NTD. **a**, Maximum likelihood phylogeny of Hsp90 protein sequences from Amorphea. The evolutionary trajectory studied, from the last common ancestor of Amorphea to modern *S. cerevisiae*, is indicated by a dark black line. Major taxonomic groups are labeled in gray. Ancestral and extant genotypes characterized in this study are in black. Complete phylogeny with taxon names is in Fig. S1. **b**, Statistical confidence in ancestral amino acid states. For each of the 98 inferred ancestral states in the NTD, the highest posterior probability of the state at any internal node along the trajectory is shown. **c**, Distribution of selection coefficients of individual ancestral states when introduced into ScHsp90, measured as the logarithm of relative fitness compared to ScHsp90 in a deep-sequencing based bulk competition assay. Dashed line indicates neutrality. Inset, close view of the region near *s* = 0. In each histogram bin, white and grey show the proportion of ancestral states with selection coefficients in that range that are estimated to be neutral or deleterious, respectively, when measurement error is taken into account.

Along this entire trajectory, substitutions occurred at 72 of the 221 sites in the NTD; because of multiple substitutions, 98 unique ancestral amino acid states existed at these sites at some point in the past and have since been replaced by the ScHsp90 state. The vast majority of these 98 ancestral states are reconstructed with high confidence (posterior probability >0.95) in one or more ancestors along the trajectory (Fig. 1b), and every ancestral sequence has a mean posterior probability across sites of >0.95 (Fig. S2a-c).

### Entrenchment and irreversibility

To measure the fitness effects of ancestral amino acids when they are re-introduced into an extant Hsp90, we created a library of ScHsp90 NTD variants, each of which contains one of the 98 ancestral states. We determined the per-generation selection coefficient (*s*) of each mutation to an ancestral state relative to ScHsp90 via bulk competition monitored by deep sequencing (Dataset S5), a technique with highly reproducible results (Fig. S3). Our assay system reduces Hsp90 expression to ~1% of the endogenous level (30), which magnifies the fitness consequences of Hsp90 mutations, enabling us to detect effects of small magnitude.

We found that the vast majority of reversions to ancestral states in ScHsp90 are deleterious (Fig. 1c). After experimental noise in fitness measurements is accounted for using a mixture model approach, an estimated 93% of all reversions reduce the fitness of ScHsp90 (95% CI 83-100%; Fig. 1c, Fig. S4). Three other statistical methods that differ in their assumptions yielded estimates that between 54% and 95% of reversions are deleterious (Fig. S5a). Two ancestral states cause very strong fitness defects (*s* = −0.38 and −0.54), but the typical reversion is only mildly deleterious (median *s* = −0.010, *P* = 1.2 × 10^−16^, Wilcoxon rank sum test; Fig. 1c). This conclusion is robust to excluding ancestral states that are reconstructed with any statistical ambiguity (*P* = 4.5 × 10^−14^). The magnitude of each mutation’s negative effect on fitness correlates with indicators of site-specific evolutionary, structural, and functional constraint, corroborating the view that they are authentically deleterious (Fig. S6).

These results do not imply that reversions can never happen—12 sites did undergo substitution and reversion at some point along the lineage from ancAmoHsp90 to ScHsp90. Rather, our observations indicate that at the current moment in time, the vast majority of ancestral states are selectively inaccessible, irrespective of whether they were available at some moment in the past or might become so in the future (31).

### Intramolecular versus intermolecular epistasis

Reversions to ancestral states might be deleterious because the derived states were entrenched by subsequent substitutions within Hsp90 (intramolecular epistasis) (1, 4); alternatively, they might be incompatible with derived states at other loci in the *S. cerevisiae* genome (intermolecular epistasis), or the derived states might unconditionally increase fitness. Entrenchment because of intramolecular epistasis predicts that introducing into ScHsp90 sets of deleterious ancestral states that existed together at ancestral nodes should not reduce fitness as drastically as would be predicted from the individual mutations’ effects. To test this possibility, we reconstructed complete ancestral NTDs from two ancestral Hsp90s separated by vast time periods on the phylogenetic trajectory (Fig. 2a, Fig. S2) and assayed their relative fitness in *S. cerevisiae* as chimeras with ScHsp90’s other domains. This design provides a lower-bound estimate of the extent of intramolecular epistasis, because it does not eliminate interactions between substitutions in the NTD and those in other domains of Hsp90.

**Figure 2.**
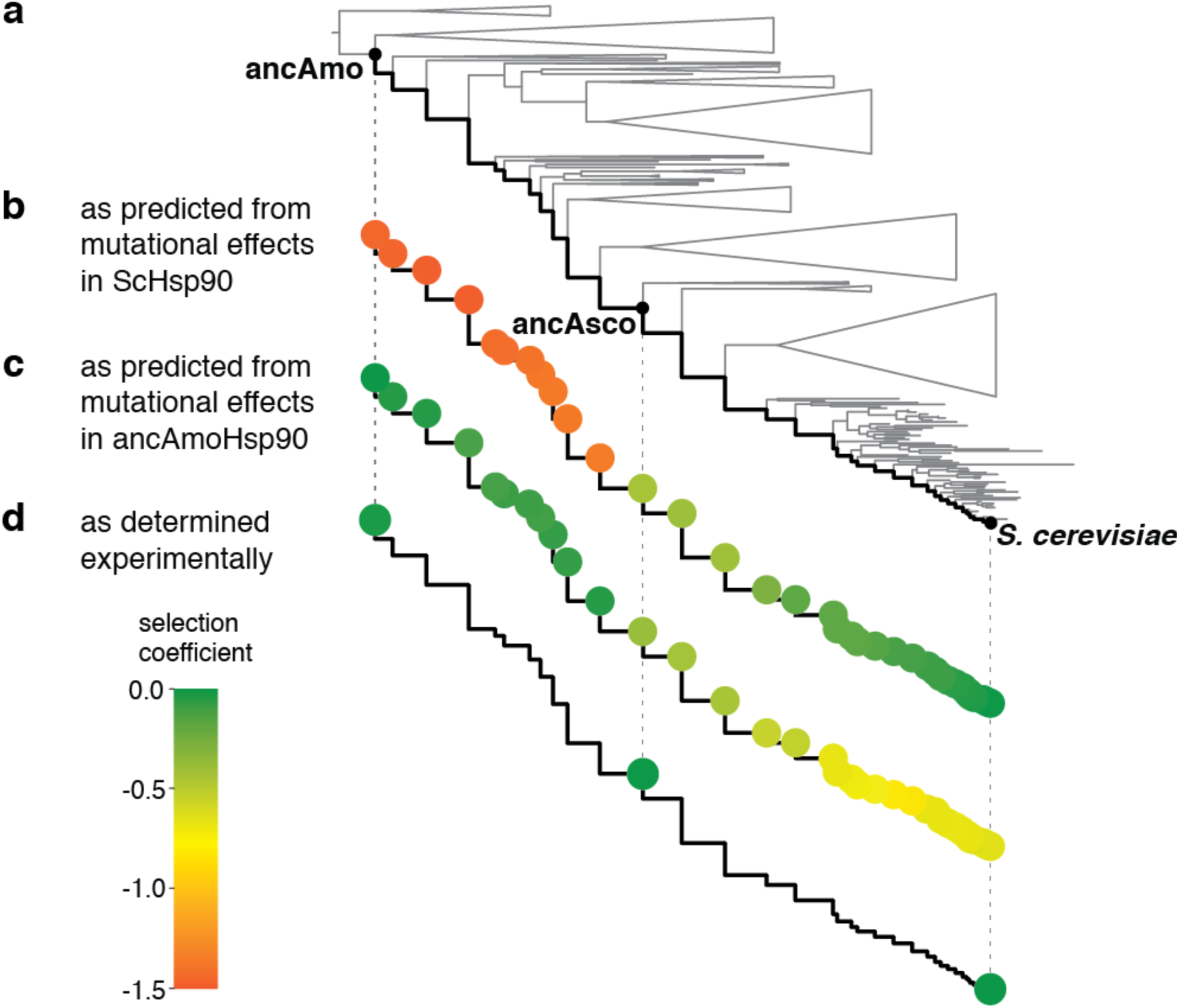
Fitness effects of historical substitutions are modified by intramolecular epistasis. For each node along the trajectory from ancAmoHsp90 to ScHsp90 (black line), the predicted or actual selection coefficient of the entire NTD genotype is represented from green (*s* = 0) to orange (*s* = −1.5). **a**, The Hsp90 phylogeny, represented as in Fig. 1a. **b**, The predicted selection coefficient of each ancestral sequence relative to ScHsp90 was calculated as the sum of the selection coefficients of each ancestral state present in that ancestor when measured individually in ScHsp90. **c**, The predicted selection coefficient of each sequence relative to ancAmoHsp90 was calculated as the sum of the selection coefficients of each derived state present at that node when measured individually in ancAmoHsp90. **d**, Experimentally determined selection coefficients for ancAmoHsp90 and ancAscoHsp90 relative to ScHsp90. For selection coefficients of each genotype, see Fig. S7.

We found that intramolecular epistasis is the predominant cause of entrenchment. The first reconstruction, ancAscoHsp90, from the ancestor of Ascomycota fungi (estimated age ~450 million years (32)), differs from ScHsp90 at 42 NTD sites. If the fitness effects of these ancestral states when combined were the same as when introduced individually, they would confer an expected fitness of 0.65 (95% CI 0.61–0.69; Fig. 2b). When introduced together, however, the actual fitness is 0.99 (Fig. 2d, Fig. S7a), indicating that the current fitness deficit of ancestral states is caused primarily by deleterious interactions within the NTD.

The older ancestor, ancAmoHsp90 (estimated age ~1 billion years (33)), differs from ScHsp90 at 60 NTD sites, which would confer an expected fitness of 0.23 (95% CI 0.21–0.26; Fig. 2b), but the actual fitness of the NTD is 0.43 (Fig. S2d, Fig. S7a), again indicating strong epistasis within the NTD. We hypothesized that the remaining fitness deficit caused by the ancAmoHsp90 NTD could be attributed to intramolecular epistasis between the NTD and substitutions in the other Hsp90 domains. We identified a candidate substitution in the protein’s middle domain that physically interacts with NTD residues to form the ATP-binding site (34); reverting this substitution to the ancAmorphea state (L378i) together with the ancAmorphea NTD increases fitness to 0.96 (Fig. 2d, Fig. S7a).

These findings indicate that virtually all the context-dependent deleterious effects of ancestral states are caused by intramolecular interactions within the NTD and with one other site in the Hsp90 protein. Derived states that emerged along the Hsp90 trajectory have been entrenched by subsequent substitutions within the same protein, which closed the direct path back to the ancestral amino acid without causing major changes in function or fitness (1).

### Contingency and permissive substitutions

We next determined whether the derived states that evolved during the protein’s history were contingent on prior permissive substitutions. We constructed a library of variants of the ancAmoHsp90 NTD, the deepest ancestor of the trajectory, each of which contains one of the 98 forward mutations to a derived state. We cloned this NTD library into yeast as a chimera with ScHsp90’s other domains (with site 378 in its ancestral state) and used our deep sequencing-based bulk fitness assay to measure the selection coefficient of each mutation relative to ancAmoHsp90 (Dataset S6, Fig. S3d,e).

We found that most mutations to derived states were selectively unfavorable (Fig. 3a). After accounting for experimental noise in fitness measurements using a mixture model, an estimated 53% of derived states reduce ancAmoHsp90 fitness (95% CI 27–96%), and 32% are neutral (95% CI 0–59%; Fig. S8); three other statistical approaches gave similar results (Fig. S5a). Fifteen percent of the derived states are beneficial in our assay (95% CI 3–57%), which could be because they are unconditionally advantageous or because of epistatic interactions with other loci in *S. cerevisiae* or other regions of ScHsp90. Two derived states had very strong fitness defects, but the typical derived state is weakly deleterious (median *s* = −0.005, *P* = 5.8 × 10^−4^, Wilcoxon rank sum test; Fig. 3a).

**Figure 3.**
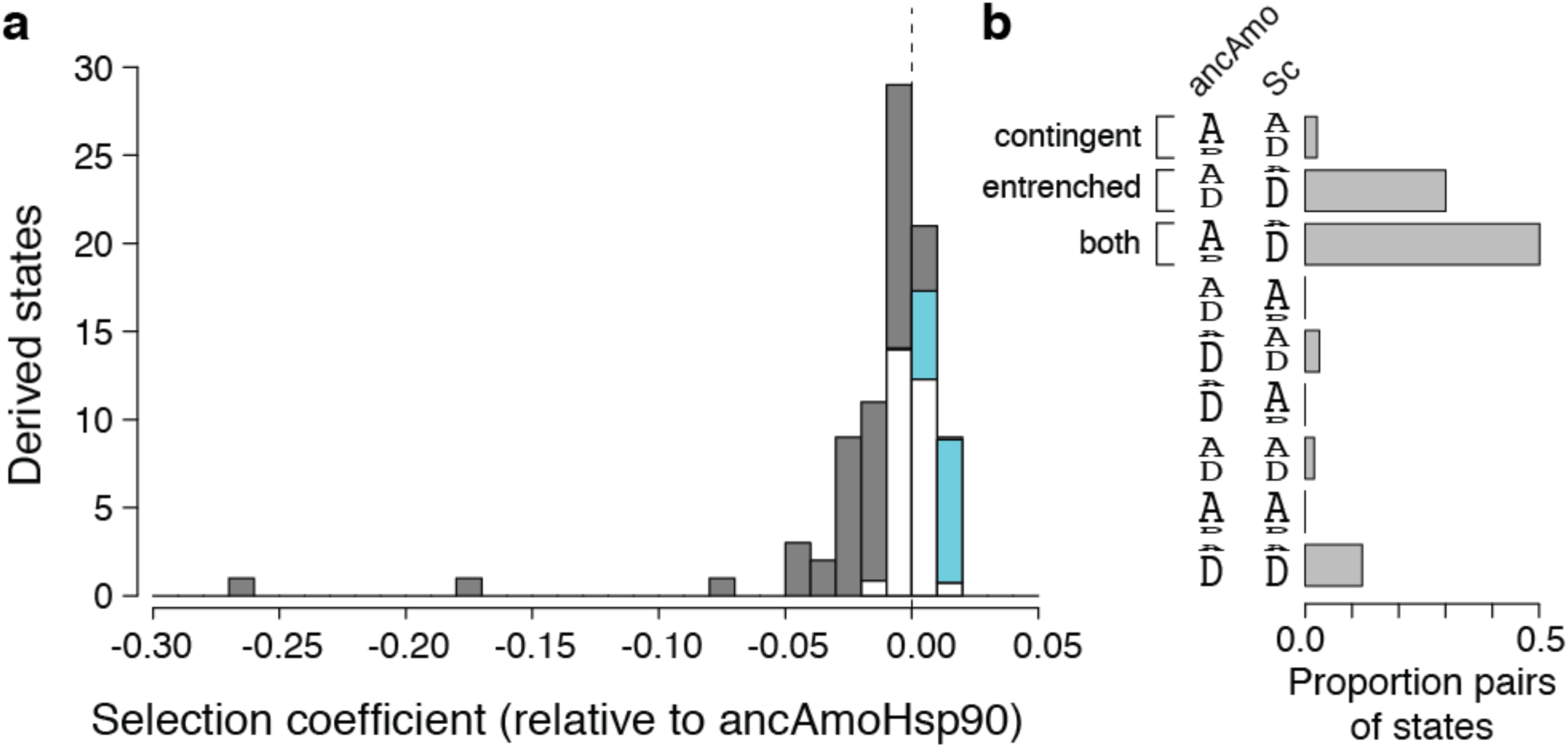
Widespread contingency and entrenchment. **a**, Distribution of measured selection coefficients of derived NTD states when introduced singly into ancAmoHsp90. Dashed line indicates neutrality. In each histogram bin, white shows the proportion of derived states with selection coefficients in that range that are estimated to be neutral; gray, deleterious; blue, beneficial. **b**, The fraction of pairs of ancestral and derived states that are inferred to be contingent, entrenched or both. Pairs of ancestral and derived states at each site can be classified by the relative fitness of the two states when measured in ancAmoHsp90 or in ScHsp90: ancestral state more fit (A larger than D), derived state more fit (D larger than A), or fitnesses indistinguishable (A and D same size). The fraction of pairs in each category was estimated as the product of the probabilities that each pair of sites is in the relevant selection category (ancestral state with fitness greater than, less than, or indistinguishable from the derived state) in the ScHsp90 and the ancAmoHsp90 backgrounds.

As with the reversions to ancestral states, the effects of individual derived states, as measured in the ancestral background, predict fitness consequences far greater than observed when the derived states are combined in the Hsp90 genotypes that existed historically along the phylogeny (Fig. 2c,d, Fig. S7b). Thus, most derived states would have been deleterious if they had occurred in the ancestral background, but they became accessible following subsequent permissive substitutions that occurred within Hsp90. Taken together, the data from the ancestral and derived libraries indicate that 83% of the amino acid states that occurred along this evolutionary trajectory were contingent on prior permissive substitutions, entrenched by subsequent restrictive substitutions, or both (Fig. 3b).

### Specificity of epistatic interactions

Epistatic effects on fitness can emerge from specific genetic interactions between substitutions that directly modify each other’s effect on some molecular property, or from nonspecific interactions between substitutions that are additive with respect to bulk molecular properties (e.g. stability (2, 35)) if those properties nonlinearly affect fitness (36-38).

To explore which type of epistasis predominates in the long-term evolution of Hsp90, we first investigated the two strongest cases of entrenchment, the strongly deleterious reversions V23f and E7a (upper-case letters indicate the ScHsp90 state and lower-case the ancestral state). We sought candidate restrictive substitutions for each of these large-effect reversions by examining patterns of phylogenetic co-occurrence. Substitution f23V occurred not only along the trajectory from ancAmoHsp90 to ScHsp90 but also in parallel on another fungal lineage; in both cases, candidate epistatic substitution i378L co-occurred on the same branch (Fig. S9a,b). As predicted if i378L entrenched f23V, we found that introducing the ancestral state i378 in ScHsp90 relieves the deleterious effect of the ancestral state f23 (Fig. 4a). These two residues directly interact in the protein’s tertiary structure to position a key residue in the ATPase active site (Fig. S9c,d), explaining their specific epistatic interaction.

**Figure 4.**
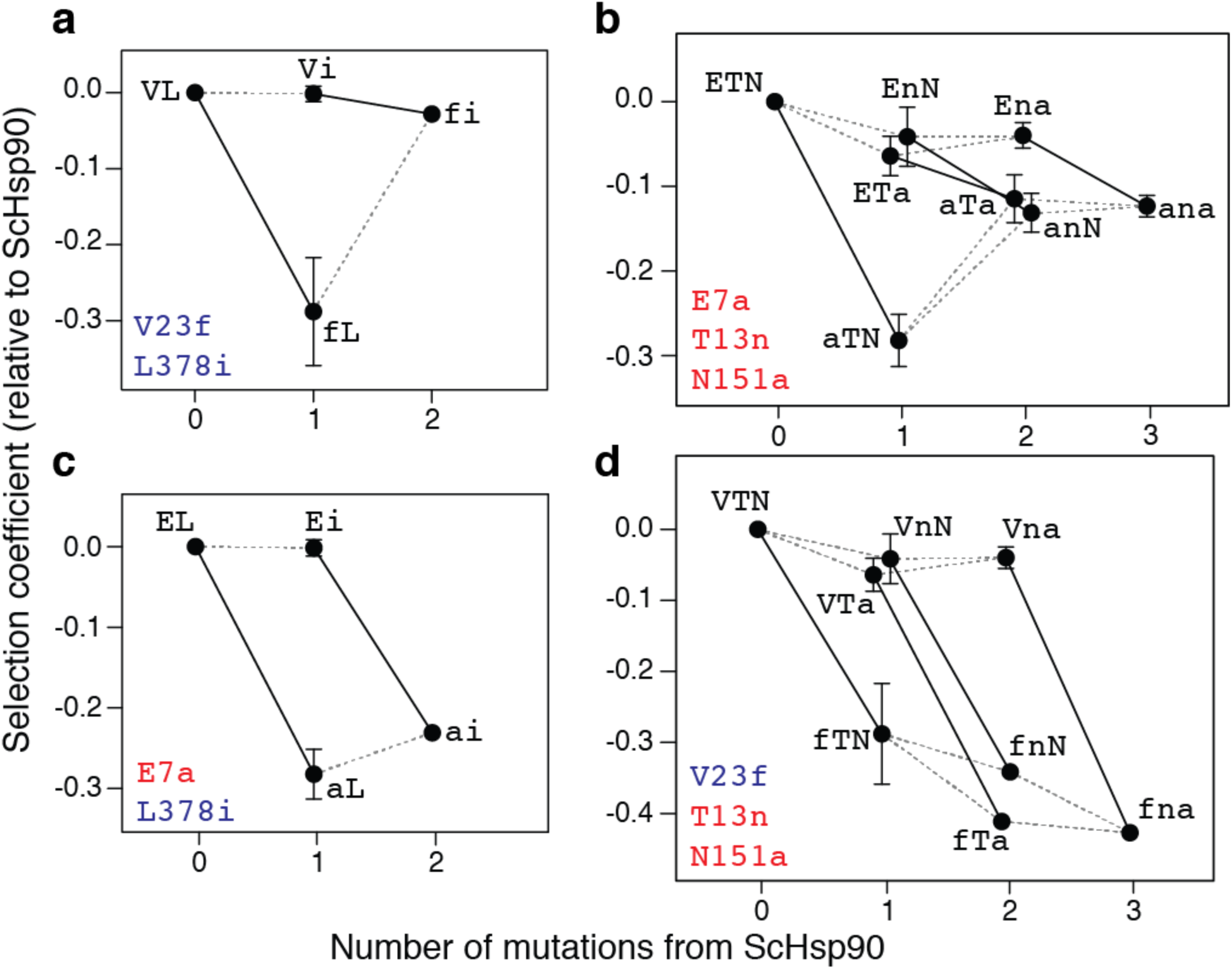
Epistatic interactions are specific. Large-effect deleterious reversions and restrictive substitutions that contributed to their irreversibility. For each single, double, or triple mutant in ScHsp90, the selection coefficient relative to ScHsp90 is shown, as assessed in monoculture growth assays. Lines connect genotypes that differ by a single mutation; solid lines indicate the effect of the large-effect reversions in each background. Error bars, SEM for 2 to 4 replicates (see Methods; absence of error bar indicates one replicate). Data points are labeled by amino acid states: lower case, ancestral state; upper case, derived state. Mutations tested in each cycle are in the bottom-left corner; those in the same color interact specifically with each other. **a**, Deleterious reversion V23f is ameliorated by L378i. **b**, Deleterious reversion E7a is partially ameliorated by N151a or T13n. **c**, L378i does not ameliorate E7a. **d**, N151a and T13n do not ameliorate V23f.

In the case of E7a—the other reversion strongly deleterious in ScHsp90—the ancestral state was reacquired in a closely related fungal lineage. We reasoned that the substitutions that entrenched a7E on the lineage leading to ScHsp90 must have themselves reverted or been further modified on the fungal branch in which reversal E7a occurred. We identified two candidates (n13T and a151N) that met these criteria (Fig. S10a,b,c). As predicted, experimentally introducing the ancestral states n13 or a151 into ScHsp90 relieves much of the fitness defect caused by the ancestral state a7, indicating that substitutions n13T and a151N entrenched a7E (Fig. 4b). These three sites are on interacting secondary structural elements that are conformationally rearranged when Hsp90 converts between ADP-and ATP-bound states (Fig. S10d,e).

To test whether these modifiers specifically restrict particular substitutions or are general epistatic modifiers, we asked whether the restrictive substitutions that entrenched one substitution also modify the effects of the other (2). As predicted if the interactions among these sets of substitutions are specific, introducing L378i does not ameliorate the fitness defect caused by E7a, and introducing T13n or N151a does not ameliorate the fitness defect caused by V23f (Fig. 4c,d). These data indicate that specific biochemical mechanisms underlie the restrictive interactions for these large-effect examples of epistatic entrenchment.

Finally, we investigated whether the epistatic interactions among the set of small-effect substitutions in this trajectory are also specific or the nonspecific result of a threshold-like relationship between fitness and some bulk property such as stability (2, 35). If epistasis is mediated by a nonspecific threshold relationship, mutations that decrease fitness in one background will never be beneficial in another, although they can be neutral if buffered by the threshold (Fig. 5a) (2, 20, 25). In contrast, specific epistatic interactions can switch the sign of a mutation’s selection coefficient in different sequence contexts (Fig. 5b) (37). As predicted under specific epistasis, we found that for most differences between ancAmoHsp90 and ScHsp90 (65%), the ancestral state confers increased fitness relative to the derived state in the ancestral background but decreases it in the extant background (Fig. 5c).

**Figure 5.**
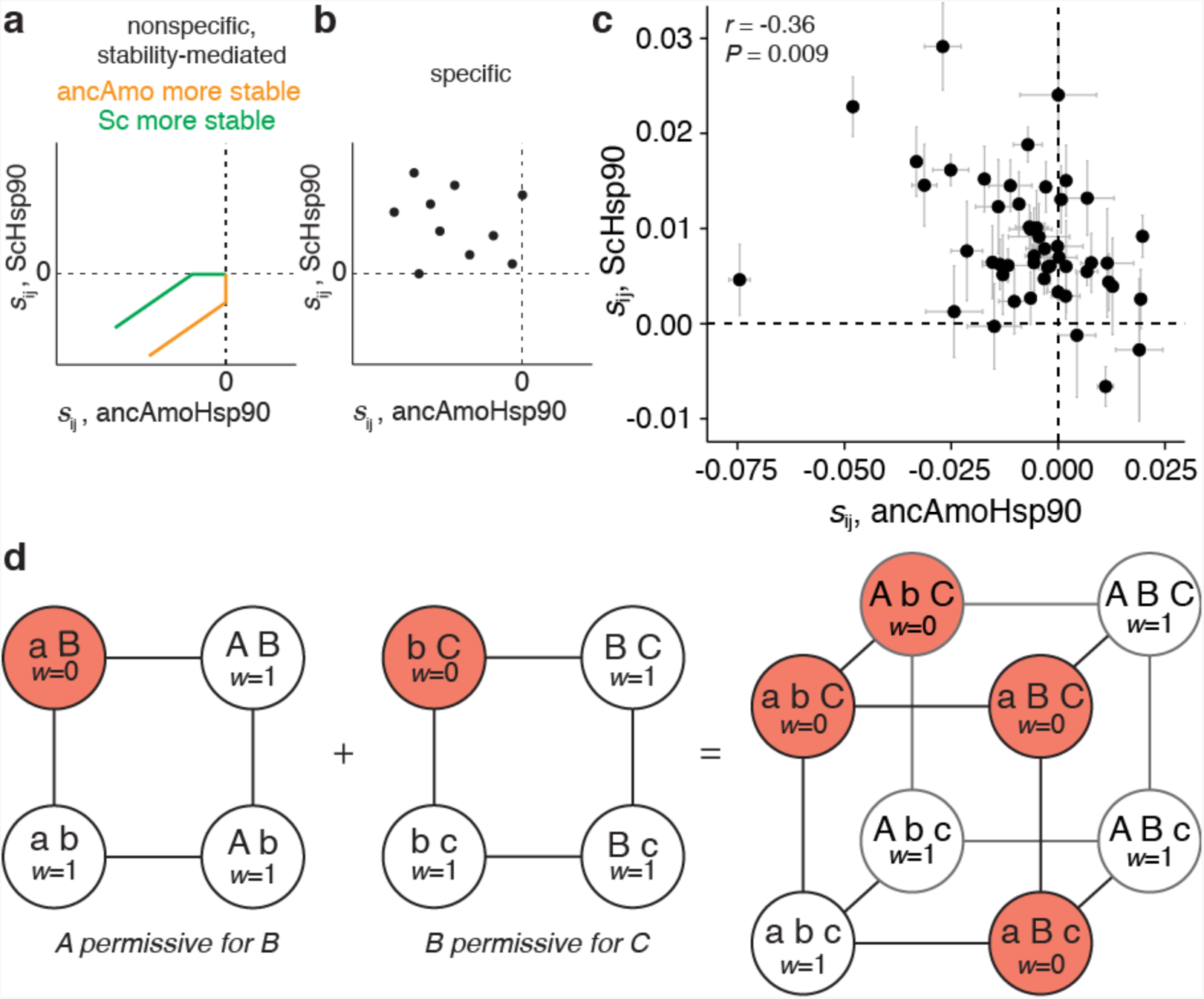
A daisy-chain model of epistasis. **a,b**, Expected relationship under two models of epistasis between selection coefficients of ancestral-to-derived mutations (*s*_ij_) when introduced into ancestral (*x*-axis) or derived (*y*-axis) backgrounds. **a**, Nonspecific epistasis: if genetic interactions are the nonspecific result of a threshold-like, buffering relationship between stability (or another bulk property) and fitness (2, 35), then the effects of strongly deleterious mutations will be positively correlated between the two backgrounds, but weakly deleterious mutations in the less stable background may be neutral in the more stable background (yellow, ancAmoHsp90 more stable; green, ScHsp90 more stable). **b**, Specific epistasis: if interactions reflect specific couplings between sites, then mutations from ancestral to derived states can be deleterious in the ancestral background but beneficial in the derived background (upper left quadrant). **c**, Measured selection coefficients for ancestral-derived state pairs that differ between ancAmoHsp90 and ScHsp90. Dashed lines, *s* = 0. Error bars, SEM from two replicate bulk competition measurements. *r*, Pearson correlation coefficient and associated *P* value. Two additional points that are strongly deleterious outliers in the ScHsp90 or ancAmoHsp90 data are not shown for clarity and are not included in the correlation; the plot including these outliers is shown in Fig. S10c. **d**, Daisy-chain model of specific epistatic interactions. Each square shows the mutant cycle for a pair of substitutions (A and B or B and C; lower-case, ancestral state; upper-case, derived), one of which is permissive for the other. Each circle is a genotype colored by its fitness (*w*): white, neutral; red, deleterious. Edges are single-site amino acid changes. The cube shows the combined mutant cycle for all three substitutions. Permissive substitutions become entrenched when the mutation that was contingent upon it occurs. Substitutions in the middle of the daisy-chain, which require a permissive mutation and are permissive for a subsequent mutation, are both contingent and entrenched.

The selection coefficients of mutations are negatively correlated between backgrounds (*P*=0.009), indicating that the substitutions that became most entrenched in the present also required the strongest permissive effect in the past. This pattern is expected if the structural constraints that determine the selective cost of having a suboptimal state at some site are conserved over time, but the specific states preferred depend on the residues present at other sites. Taken together, these findings indicate that most epistasis during the long-term evolution of the Hsp90 NTD involved specific one-to-one (or few-to-few) interactions among sites, not general effects on the protein’s tolerance to mutation.

## Discussion

### Relation to prior work

We observed widespread and specific epistasis over the course of a billion years of Hsp90 evolution, during which the protein’s function, physical architecture, and fitness were conserved. The fraction of historical substitutions that were either contingent on permissive substitutions or entrenched by restrictive substitutions—about 80%—is considerably higher than suggested by previous experimental work (2, 22, 24) and some computational analyses (14), rivaling the highest estimates from computational studies (1, 16, 17). One explanation for the more widespread epistatic interactions in our study may be our method’s capacity to detect much smaller growth deficits than have been discernable in previous experimental studies.

Another difference from previous research is that we primarily observed specific epistasis, whereas several studies have found a dominant role for nonspecific stability-mediated epistasis, particularly during the short-term evolution of viruses (2, 11, 20). This disparity could be attributable to a difference in selective regime or in time scale: the epistatic constraints caused by specific interactions are expected to be maintained over far longer periods of time than those caused by nonspecific interactions, which are easily replaced by other substitutions because of the many-to-many relationship between permissive and permitted amino acid states (2, 20, 37). The prevalence and type of epistasis may also vary because of differences in proteins’ physical architectures. Additional case studies will be necessary to evaluate the causal role of these and other factors in determining the nature of epistatic interactions during evolution.

### Limitations

Our strategy has some known limitations, but none are likely to change our major conclusions. For example, we assayed the effect of long-past substitutions in the context of extant yeast cells. Our experiments, however, indicate that there is only very weak epistasis for fitness between historical substitutions within Hsp90 and those at other loci, because the reconstructed ancestral Hsp90 chimeras cause a fitness deficit of only 0.01 to 0.04 when introduced into *S. cerevisiae* cells—much smaller than the sum of intramolecular incompatibilities revealed by introducing the ancestral states individually. Our finding of widespread contingency and entrenchment is therefore not an artifact of incompatibilities between ancestral Hsp90 states and the genotype of present-day *S. cerevisiae* at other loci.

A second potential limitation is that the ancestral states we tested were reconstructed phylogenetically, not known empirically. But the vast majority of states were inferred with high statistical confidence, because the Hsp90 NTD is well conserved and we used a densely-sampled alignment. The ambiguity that was present primarily concerned the specific ancestral node at which an inferred ancestral state was present, not whether or not it was ancestral somewhere along the trajectory, which is the key inference for our purposes. Further, even when all states with any degree of statistical uncertainty in the ancestral reconstruction were excluded from the analysis, the remaining data strongly supported our conclusions concerning contingency and entrenchment.

Finally, we measured fitness under a particular set of experimental conditions. Our assay system reduces Hsp90 expression to ~1% of the endogenous level (30). Based on previous work quantifying the relationship between Hsp90 function, expression, and growth rate (30), we estimate that the average selection coefficient of −0.01 we observed among contingent or entrenched substitutions corresponds to a fitness deficit of approximately *s* = −5 × 10^−6^ under native-like expression levels. Mutations with selection coefficients in this range would likely be subject to purifying selection in large microbial populations (39-41). Our assay also tests fitness under log-phase growth conditions in rich media. A more heterogeneous or demanding environment would likely increase the magnitude of selective effects of Hsp90 mutations, because stress should amplify the fitness consequences of mutations in the proteostasis machinery.

### Implications

Our observation that contingency and entrenchment affected the majority of historical substitutions suggests a daisy-chain model by which genetic interactions structured long-term Hsp90 evolution (Fig. 5d). A permissive mutation becomes entrenched and irreversible once a substitution contingent upon it occurs; if the contingent substitution subsequently permits a third substitution, it too becomes entrenched (1, 16).

Most of the substitutions along the trajectory from ancAmoHsp90 to ScHsp90 were both contingent and entrenched, suggesting that they occupy an internal position in this daisy chain. Each of these changes closed reverse paths at some sites and opened forward paths at others, which—if taken—would then entrench the previous step. Evolving this way over long periods of time, proteins come to appear exquisitely well-adapted to the conditions of their existence, with most present states superior to past ones. The conditions that make today’s states so fit, however, include—or are even dominated by—the transient internal organization of the protein itself.

## Methods

For additional details, see SI Methods.

### Phylogenetic inference and ancestral reconstruction

We inferred the maximum likelihood (ML) phylogeny for 261 Hsp90 protein sequences from the Amorphea clade, with Viridiplantae as an outgroup, under the LG+Γ+F model in RAxML version 8.1.17 (42). Most probable ancestral NTD sequences were reconstructed on this ML phylogeny using the AAML module of PAML version 4.4 (43). The trajectory of sequence change was enumerated from the amino acid sequence differences between successive ancestral nodes on the lineage from the common ancestor of Amorphea (ancAmoHsp90) to *S. cerevisiae* Hsp82 (ScHsp90). Ancestral states are defined as amino acid states not present in ScHsp90 that occurred in at least one ancestral node on the lineage from ancAmoHsp90 to ScHsp90. Derived states are defined as amino acid states not present in the reconstructed ancAmoHsp90 sequence that occurred in at least one descendent node on the lineage to ScHsp90.

### Bulk growth competitions

The ScHsp90 and ancAmoHsp90 NTD (+L378i, see SI Methods, Fig S2d) protein-coding sequences were expressed together with the other domains from ScHsp90 from the p414ADHΔTer plasmid (30). Individual mutations in each variant library were introduced via PCR. Library genotypes were tagged with short barcodes to simplify sequencing steps during the bulk competition (44); barcodes were associated with variant genotypes via paired-end sequencing on an Illumina MiSeq instrument. Variant libraries were transformed into the DBY288 Hsp90 *S. cerevisiae* shutoff strain (45, 46) and grown for 48 (ScHsp90 library) or 31 (ancAmoHsp90 library) hours under selective conditions. Cultures were maintained in log-phase by regular dilution with fresh media to maintain a population size of 10^9^ or greater, and samples of ~10^8^ cells were collected at regular time points over the course of the bulk competition. Plasmid DNA was isolated from each time point (47), and the frequency of each library genotype at each time point was determined via Illumina sequencing. Bulk competitions were performed in duplicate.

### Determination of selection coefficients

The ratio of the frequency of each variant in the library relative to wildtype (ancAmoHsp90 or ScHsp90) was determined from the number of sequence reads at each time point, and the slope of the logarithm of this ratio versus time (in number of generations) was determined as the raw per-generation selection coefficient (*s*) (48):

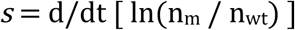

where n_m_ and n_wt_ are the number of sequence reads of mutant and wildtype, respectively, and time is measured in number of wildtype generations. For genotypes that were not assayed in the bulk competition, selection coefficients were determined from monoculture growth rates (48), measured over 30 hours of growth with periodic dilution to maintain log-phase growth (30). Selection coefficients for both types of measurement were scaled in relative fitness space (*w* = e^*s*^) such that an Hsp90 null allele, which is lethal, has a relative fitness of 0 (*s* = -∞).

### Estimating the fraction of deleterious mutations

We estimated the fraction of mutations in each library that are deleterious by fitting the distribution of mutant selection coefficients to a mixture model of underlying Gaussian distributions. One of the underlying mixture components was required to have the mean and standard deviation derived from replicate measurements of the wildtype sequences that were present in the library but represented by independent barcodes; remaining mixture components had a freely fit mean and standard deviation, and all components had a freely fit mixture proportion. The optimal number of mixture components was determined via Akaike Information Criterion. The mixture component derived from the wildtype sampling distribution was taken to represent genotypes in the library with fitness indistinguishable from wildtype; mixture components with mean less than zero were taken to reflect deleterious mutations; and mixture components with mean greater than zero were taken to reflect beneficial mutations. For each variant, the posterior probability of being neutral, deleterious, or beneficial was determined from the relative probability density function for mixture components in each category at the selection coefficient measured for that mutation; the total fraction of variants in the library that are deleterious (or beneficial) was determined by summing the posterior probabilities of being deleterious (or beneficial) over all mutants.

To determine the robustness of the mixture model’s estimates of the fraction of deleterious (or beneficial) mutations, we used three other approaches to analyze the distribution of fitness measurements. The simplest, a nonparametric approach, reports the proportion of mutations with observed selection coefficients < 0; because error in fitness measurements appears to be unbiased (Fig. S5) and the number of deleterious mutations exceeds the number of beneficial mutations, this approach is expected to yield a slightly conservative estimate of the true proportion of deleterious mutations with selection coefficients of any magnitude. The second approach is an empirical Bayes approach that calculates the posterior probability that each mutation is deleterious (or beneficial), given the experimentally observed selection coefficients and measurement error for wildtype and mutant genotypes; these posterior probabilities are summed to yield the estimated proportion of mutations in each fitness category. Third, we constructed 95% confidence intervals for each mutation’s selection coefficient given its mean over two replicates and standard error estimated over all mutations and counted the number of mutations with selection coefficients less (or greater) than zero whose confidence intervals do not overlap zero.

### Data and code availability

Processed sequencing data and scripts to reproduce all analyses are available at github.com/JoeThorntonLab/Hsp90_contingency-entrenchment. Tables listing mutants and their selection coefficients are included as Datasets S5 and S6. Raw sequencing data from each bulk competition have been deposited in the NCBI Sequence Read Archive under accession SRP126524. Tables linking barcode variants to their associated Hsp90 genotype are included as Datasets S7 and S8.

## Acknowledgements

We thank members of the Thornton lab for comments on the manuscript and Jeff Boucher for technical assistance. This work was supported by National Institutes of Health R01GM104397 and R01GM121931 (J.W.T.), R01GM112844 (D.N.A.B.), F32GM119205-2 (J.M.F.), T32-GM007183 (T.N.S.), Government of India Department of Biotechnology Ramalingaswami Fellowship (P.M.), and a National Science Foundation Graduate Research Fellowship (T.N.S.)

## Author Contributions

All authors conceived the project and designed experiments. T.N.S., J.M.F., and P.M. performed experiments and analyzed data. T.N.S. and J.W.T. wrote the paper, with contributions from all authors.

## Author Information

The authors declare no competing financial interests. Correspondence and requests for materials should be addressed to J.W.T. or D.N.A.B.

## Supporting Information

### SI Methods

#### Phylogenetic analysis and ancestral reconstruction

We obtained Hsp90 protein sequences from the Amorphea clade (29) from NCBI, the JGI Fungal Program, the Broad Institute Multicellularity Project, the literature (49), and Iñaki Ruiz-Trello (personal communication). Full identifiers and sources of sequences are listed in Dataset S3. Each protein was used as a query in a BLASTp search against the human proteome to identify and retain Hsp90A orthologs. We used CD-HIT (50) to filter proteins with high sequence similarity. We removed sequences with >67% missing characters and highly diverged, unalignable sequences. Remaining sequences were aligned with Clustal Omega (51). Lineage-specific insertions were removed, as were unalignable linker regions (ScHsp90 sites 1-3, 225-237, 686-701). We added six Hsp90A sequences from Viridiplantae as an outgroup, resulting in a final alignment of 267 protein sequences and 680 sites (Dataset S1).

We inferred the maximum likelihood (ML) phylogeny (Dataset S2) given our alignment and the LG model (52) with gamma-distributed among-site rate variation (4 categories) and ML estimates of amino acid frequencies, which was the best-fit model as judged by AIC. The phylogeny was inferred using RAxML version 8.1.17 (42). The ML phylogeny reproduces accepted relationships between major taxonomic lineages (29, 53-57). Most probable ancestral sequences (Dataset S4) were reconstructed on the maximum likelihood phylogeny using the AAML module of PAML version 4.4 (43) given the alignment, ML phylogeny, and LG+G model. The trajectory of sequence change was enumerated from the amino acid sequence differences between successive ancestral nodes on the lineage from the common ancestor of Amoebozoa + Opisthokonta (ancAmorphea) to *S. cerevisiae* Hsp82 (ScHsp90, Uniprot P02829).

Coding sequences for the most probable ancestral amino acid sequences of the Hsp90 N-terminal domain (NTD) from ancAmorphea (ancAmoHsp90) and the common ancestor of Ascomycota yeast (ancAscoHsp90) were synthesized by IDT (Dataset S9). These sequences were cloned as chimeras with the ScHsp90 middle and C-terminal domains and intervening linkers via Gibson Assembly. AncAmoHsp90 also carries an additional reversion to the ancAmorphea state at site 378 in the middle domain (Fig. S2d), which is part of a loop that extends down and interacts with ATP and the NTD (34, 58).

#### Generating mutant libraries

ScHsp90 and ancAmoHsp90 gene constructs were expressed from the p414ADHΔTer plasmid (30). The ScHsp90 library consists of variants of the ScHsp90 NTD, each containing one mutation to an ancestral amino acid state. The ancAmoHsp90 library consists of variants of the ancAmoHsp90 NTD, each containing one mutation to a derived state. Two sets of PCR primers were designed for each mutation, to amplify Hsp90 NTD fragments N-terminal and C-terminal to the mutation of interest; primers introduce the mutation of interest and generate a 25-bp overlap between fragments, as well as 20-bp overlaps between each fragment and the destination vector for gene re-assembly (Dataset S9). PCR was conducted with Pfu Turbo polymerase (Agilent) for 15 amplification cycles. The resulting PCR fragments were stitched together with a 10-cycle assembly PCR, pooled, and combined via Gibson Assembly (NEB) with a linearized p414ADHΔTer Hsp90 destination vector excised of the NTD.

#### Barcode labeling of library genotypes

Following construction of the plasmid libraries, each variant in the library was tagged with a unique barcode to simplify sequencing steps during bulk competition (44). A pool of DNA constructs containing a randomized 18 base-pair barcode sequence (N18) and Illumina sequencing primer annealing regions (IDT; Dataset S9) was cloned 200 nucleotides downstream from the hsp90 stop codon via restriction digestion, ligation, and transformation into chemically-competent *E. coli*. Cultures with different amounts of the transformation reaction were grown overnight and the colony forming units in each culture were assessed by plating a small fraction. We isolated DNA from the transformation that contained approximately 10-20 fold more colony-forming units than mutants, with the goal that each mutant would be represented by 10-20 unique barcodes.

To associate barcodes with Hsp90 mutant alleles, we conducted paired end sequencing of each library using primers that read the N18 barcode in the first read and the Hsp90 NTD in the other (Dataset S9). To generate short DNA fragments from the plasmid library that would be efficiently sequenced, we excised the gene region between the NTD and the N18 barcode via restriction digest, followed by blunt ending with T4 DNA polymerase (NEB) and plasmid ligation at a low concentration (3 ng/μL) that favors circularization over bi-molecular ligations. The resulting DNA was re-linearized by restriction digest, and Illumina adapter sequences were added via an 11-cycle PCR (Dataset S9). The resulting PCR products were sequenced using an Illumina MiSeq instrument with asymmetric reads of 50 bases and 250 bases for Read1 and Read2 respectively. After filtering low quality reads (Phred scores < 10), the data were organized by barcode sequence. For each barcode that was read more than 3 times, we generated a consensus sequence of the N-domain indicating the mutation that it contained (Datasets S7, S8).

#### Bulk growth competitions

For bulk fitness assessments, we transformed *S. cerevisiae* with the ScHsp90 library along with wildtype ScHsp90 and a no-insert control; we also transformed *S. cerevisiae* with the ancAmoHsp90 library along with wildtype ScHsp90, wildtype ancAmoHsp90, and a no-insert control. Concentrations of plasmids were adjusted to yield a 2:6:1 molar ratio of wildtype: no-insert control: average variant in the library. Plasmid libraries and corresponding controls were transformed into the DBY288 Hsp90 shutoff strain (45, 46), resulting in ~150,000 unique yeast transformants representing 50-fold sampling for the average barcode. Following recovery, transformed cells were washed 5 times in SRGal-W (synthetic 1% raffinose and 1% galactose lacking tryptophan) media to remove extracellular DNA, and then transferred to plasmid selection media SRGal-W and grown at 30°C for 48 hours with repeated dilution to maintain the cells in log phase of growth. To select for function of the plasmid-borne Hsp90 allele, cells were shifted to shutoff conditions by centrifugation, washing and re-suspension in 200 mL SD-W (synthetic 2% dextrose lacking tryptophan) media and ampicillin (50μg/mL), and growth at 30°C 225 rpm. Following a 16-hour growth period required to shut off expression of the wildtype chromosomal Hsp90, we collected samples of ~10^8^ cells at 8 or more time points over the course of 48 (ScHsp90 library) or 31 (ancAmoHsp90 library) hours and stored them at −80°C. Cultures were maintained in log phase by regular dilution with fresh media, maintaining a population size of 10^9^ or greater throughout the bulk competition. Bulk competitions of each library were conducted in duplicate from independent transformations.

#### DNA preparation and sequencing

We collected plasmid DNA from each bulk competition time point as previously reported (47). Purified plasmid was linearized with AscI. Barcodes were amplified by 18 cycles of PCR using Phusion polymerase and primers that add Illumina adapter sequences, as well as an 8-bp identifier used to distinguish among libraries and time points (Dataset S9). Identifiers were designed so that each differed by more than two bases from all others to avoid misattributions due to sequencing errors. PCR products were purified two times over silica columns (Zymo research), and quantified using the KAPA SYBR FAST qPCR Master Mix (Kapa Biosystems) on a Bio-Rad CFX machine. Samples were pooled and sequenced on an Illumina NextSeq (ancAmoHsp90 library) or HiSeq 2000 (ScHsp90 library) instrument in single-end 100 bp mode.

#### Analysis of bulk competition sequencing data

Illumina sequence reads were filtered for Phred scores >20, strict matching of the sequence of the intervening bases to the template, and strict matching of the N18 barcode and experimental identifier to those that were expected in the given library. Reads that passed these filters were parsed based on the identifier sequence. For each identifier, the data was condensed by generating a count of each unique N18 read. The unique N18 count file was then used to identify the frequency of each mutant using the variant-barcode association table. For each variant in the library, the counts of each associated barcode were summed to generate a cumulative count for that mutant.

#### Determination of selection coefficient

The ratio of the frequency of each variant in the library relative to wildtype (ancAmoHsp90 or ScHsp90) was determined at each time point, and the slope of the logarithm of this ratio versus time (in number of generations) was determined as the raw per-generation selection coefficient (*s*) (48):

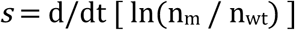

where n_m_ and n_wt_ are the number of sequence reads of mutant and wildtype, respectively, and time is measured in number of wildtype generations. No-insert plasmid selection coefficients were determined from the first three time points because their counts drop rapidly over time. Mutants with selection coefficients within three standard deviations of the mean of no-insert variants were considered null-like and also analyzed based on the first three time points. For all other variants, selection coefficients were determined from all time points. Final selection coefficients for each variant were scaled in relative fitness space (*w* = e^*s*^) such that the Hsp90 null allele, which is lethal, has a relative fitness of 0 (*s* = -∞). This definition of relative fitness, unlike that which defines *w* = 1 – *s*, has the advantage of making selection coefficients additive and reversible (the selection coefficient of mutation from state *i* to *j* is the opposite of the selection coefficient of that from *j* to *i*) (59).

#### Generation of individual mutants and monoculture analysis of yeast growth

To measure the relative fitness of ancAscoHsp90, mutations missed in the bulk libraries, and genotypes in mutant cycles that we sought to test in combination for epistatic interactions, we assayed growth rate in monoculture and related this to fitness, which assumes the relative rate of growth of two genotypes is the same in isolation as in direct competition (48). The growth rate of individually cloned mutants was estimated over 30 hours of growth with periodic dilution to maintain log-phase growth, as per Jiang et al. (30). Growth rates were determined as the slope of the linear model relating the log-transformed dilution-corrected cell density to time. The growth rate was converted to an estimate of the selection coefficient by taking the difference in growth rate (Malthusian parameter) between mutant and wildtype and multiplying this by the wildtype generation time (48), then rescaling selection coefficients in relative fitness space such that a null mutant has relative fitness 0 (*s* = -∞).

Individual mutants of ancAmoHsp90 and ScHsp90 were generated in the p414ADHΔTer background by Quikchange site-directed mutagenesis (Dataset S9), confirmed by Sanger sequencing. Mutations that were generated and assayed in ancAmoHsp90 (with number of replicate measurements in parentheses) include: S49A (n=1), T137I (n=1), V147I (n=1), I158V (n=1), R160L (n=1), G164N (n=1), E165P (n=1), L167I (n=1), K172I (n=1), L193I (n=1), and V194I (n=1). Mutations generated and assayed in ScHsp90 include: T5S (n=3), E7A (n=4), T13N (n=3), V23F (n=2), N151A (n=3), L378I (n=2), double mutants E7A/T13N (n=3), E7A/N151A (n=3), T13N/N151A (n=3), V23F/T13N (n=1), V23F/N151A (n=1), E7A/L378I (n=1), V23F/L378I (n=1) and triple mutants E7A/T13N/N151A (n=2) and V23F/T13N/N151A (n=1).

#### Robustness of results to statistical uncertainty and technical variables

The conclusion that the typical ancestral state is deleterious in ScHsp90 is robust to the exclusion of 20 ancestral states that have posterior probability < 1.0 at all ancestral nodes along the trajectory (*P* = 4.5 × 10^−14^, Wilcoxon rank sum test with continuity correction). The mutation to one ancestral state was missed in the bulk competition: its selection coefficient was inferred separately via monoculture, and including it in the analysis still leads to the conclusion that the typical ancestral state is deleterious (*P* = 7.8 × 10^−17^, Wilcoxon rank sum test with continuity correction).

The conclusion that the average derived state is deleterious in ancAmoHsp90 is retained when we include only the 32 mutations for which the ancAmoHsp90 state is inferred with a posterior probability of 1.0 and the derived state is inferred with posterior probability 1.0 in at least one node along the trajectory (*P* = 1.1 × 10^−4^, Wilcoxon rank sum test with continuity correction). The conclusion is also robust if we include selection coefficients as determined separately via monoculture for mutations to 11 derived states that were missed in the bulk competition (*P* = 5.4 × 10^−4^, Wilcoxon rank sum test with continuity correction).

We assessed relative fitness for six genotypes (ScHsp90+E7a, ScHsp90+V23f, ScHsp90+N151a, ScHsp90+T13n, ancAmoHsp90, and ancAmoHsp90+i378L) both by monoculture and by bulk competition. These two measures are well correlated (Pearson R^2^ = 0.95), although the magnitude of a fitness effect is smaller when measured by monoculture growth assays (Fig. S3f), perhaps because of differences in experimental conditions for bulk versus monoculture growth, such as the type of growth vessel and culture volume (and consequential aeration). The only conclusion involving a comparison between these two kinds of measurements is that ancAscoHsp90 (measured via monoculture) is more fit than would be predicted from the sum of selection coefficients of its component states (measured via bulk competition) (Fig. 2, Fig. S7). We therefore used the observed linear relationship between the two types of fitness assays to transform ancAscoHsp90’s fitness as measured by monoculture (0.991); the expected fitness of ancAscoHsp90 in a bulk competition is 0.986, still much larger than the predicted fitness of 0.65 in the absence of epistasis.

#### Expected versus observed fitness

To identify epistasis between candidate interacting sites (e.g. Fig. 4a-d) or among the broader set of substitutions (e.g. Fig. 2), we compared the observed fitness of genotypes with multiple mutations to that expected in the absence of epistasis. In the absence of epistatic interactions, selection coefficients combine additively (59). We therefore calculated the expected selection coefficient of a genotype as the sum of selection coefficients of its component mutations as measured independently in a reference background (ancAmoHsp90 or ScHsp90). The standard error of a predicted fitness given the sum of selection coefficients was calculated as the square root of the sum of squared standard errors of the individual selection coefficient estimates, as determined from the duplicate bulk competition measurements. Epistasis was implicated if the observed fitness of a genotype differed from that predicted from the sum of its corresponding single-mutant selection coefficients.

#### Estimating the fraction of deleterious mutations

We sought to determine the fraction of mutations in each dataset that are deleterious using a modeling approach that incorporates measurement error and which does not require individual mutations to be classified as deleterious, neutral, or beneficial. We used the mixtools package (60) in R to estimate mixture models of underlying Gaussian distributions that best fit the observed distributions of mutant selection coefficients in each library. First, we fit a single Gaussian distribution to the measured selection coefficients of replicate wildtype sequences that were present in the library but represented by independent barcodes. We then required one of the Gaussian distributions in each mixture model to have a mean and standard deviation fixed to that of the wildtype measurements, with a freely estimated mixture proportion. The other Gaussian components in each mixture model had a freely fit mean, standard deviation, and mixture proportion. Mixture models were fit to all non-outlier selection coefficients, because the presence of strongly deleterious selection coefficients (*s* < −0.04), which are unambiguously deleterious, interfered with model convergence. We assessed mixture models with a variable number of mixture components (*k* = 2 to 6 for the ancAmoHsp90 library and 2 to 5 for the ScHsp90 library, because the 6-component model would not converge), and obtained the maximum likelihood estimate of each component’s mean, standard deviation, and mixture proportion via an expectation-maximization algorithm as implemented in mixtools. We compared the models built for each *k* using AIC. For ScHsp90, the 3-component mixture model was favored by AIC (Fig. S4a). For ancAmoHsp90, the 2-component and 5-component mixture models had virtually indistinguishable AIC (Fig. S8a), but the 2-component mixture model had a visually suboptimal fit (Fig. S8c,d) and attributed a larger proportion of mutations as belonging to a deleterious sampling distribution (0.78 versus 0.53 for the 5-component mixture model), so we selected the more conservative and visually superior 5-component mixture model.

The mixture component derived from the wildtype sampling distribution was taken to represent genotypes in the library with fitness indistinguishable from wildtype; mixture components with mean < 0 were taken to reflect deleterious variants; and mixture components with mean > 0 were taken to reflect beneficial variants. For each variant, the posterior probability of being deleterious, neutral, or beneficial was determined from the relative probability density function for mixture components in each category at the selection coefficient measured for that mutation; for variants with *s* < −0.04 that were excluded from model inference, the posterior probability of being deleterious was 1. The total fraction of variants in the library that are deleterious (or beneficial) was determined by summing the posterior probabilities of being deleterious (or beneficial) over all mutants. To generate the representations in Figures 1c and 3a, posterior probabilities were summed separately for the set of measurements that fall within each histogram bin. We also report the estimate of the fraction deleterious (or beneficial) by summing the mixture proportion parameters for mixture components centered below (or above) zero (Figs. S4a, S8a).

Uncertainty in the estimated fraction of mutations that are deleterious or beneficial was determined via a bootstrapping procedure. For each of 10,000 bootstrap replicates, measured selection coefficients from the bulk competition were resampled with replacement. Mixture models with fixed *k* were fit to each bootstrap sample, and the estimated fractions of mutations in deleterious, neutral, or beneficial sampling distributions were determined as above.

To estimate the probability that a pair of states exhibit contingency and/or entrenchment, we calculated the joint posterior probability as the product of the probabilities that each pair of sites is in the relevant selection category (ancestral state with fitness greater than, less than, or indistinguishable from the derived state) in the ScHsp90 and the ancAmoHsp90 backgrounds. For sites that substituted from the ancAmoHsp90 state *i* to the ScHsp90 state *j* (*i*→*j*, n = 35), *i* is the ancestral state and *j* the derived state for measurements in both backgrounds. For sites that substituted from the ancAmoHsp90 state *i* to an intermediate state *j* before substituting back to *i* in ScHsp90 (*i*→*j*→*i*, n=12), then *i* is the ancestral state and *j* derived in ancAmoHsp90 assay, and *j* is the ancestral state and *i* derived in ScHsp90. For sites that substituted from the ancAmoHsp90 state *i* to an intermediate state *j* that was further modified to *k* in ScHsp90 (*i*→*j*→*k*, n=25), two comparisons were made: in the first, *i* was ancestral and *k* was derived for measurements in both backgrounds, while in the second comparison, *i* was ancestral and *j* derived in ancAmoHsp90, and *j* ancestral and *k* derived in ScHsp90.

In addition to the mixture model approach presented above, we report three independent methods for estimating the fraction of mutations in each dataset that are deleterious (or beneficial) (Fig. S5a). The simplest estimate of the fraction of mutations in each distribution that are deleterious (or beneficial) is the fraction of observed selection coefficients (*s*_obs_) that are less than (or greater than) zero. This counting approach assumes that, at some magnitude, all mutations have a true *s* > 0 or *s* < 0. This method would be unbiased if experimental errors are random and if the number of truly beneficial and truly deleterious mutations is equal. In our data, experimental errors are unbiased with respect to *s*_obs_ (Fig. S5b,c), but there appear to be more deleterious than beneficial mutations. As a result, measurement error is likely to cause the number of mutations with true *s* < 0 and *s*_*ob s*_> 0 to exceed the number with true *s* > 0 and *s*_*obs*_ < 0; this approach is therefore expected to underestimate the fraction of mutations with true *s* < 0.

Second, we used an empirical Bayes approach. For each mutation, we compute the posterior probability that it is non-neutral by comparing the likelihoods of two hypotheses: the null hypothesis, that a variant is neutral and therefore *s* ~ N(0, SEM_wt_) where SEM_wt_ is the standard deviation of the sampling distribution of repeated wildtype measurements present in the corresponding experiment (Figs. S4b, S8b); and the alternative hypothesis, that a variant is non-neutral and therefore *s* ~ N(*s*_obs_, SEM_mut_), where SEM_mut_ is calculated as an estimated SEM from all duplicate bulk fitness measurements, which makes the assumption that all variants have the same experimental error (Fig. S5b,c). SEM_mut_ was calculated as:

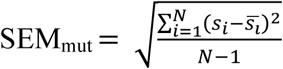

where *s*_*i*_ is a measured selection coefficient of a mutant in a single replicate, 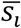 is the corresponding mean selection coefficient for that mutant as calculated from both replicates, and *N* is the total number of observations from both replicates. The posterior probability that a variant is non-neutral is calculated from the relative likelihoods of the two hypotheses, with a uniform prior on the two hypotheses:

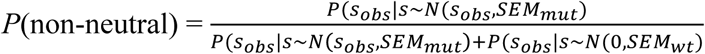

If a variant has *s*_obs_ > 0, then *P*(non-neutral) corresponds to a probability that a mutation is beneficial; if a variant has *s*_obs_ < 0, then *P*(non-neutral) corresponds to a probability that a mutant is deleterious. Like the counting approach, this empirical Bates approach will call some truly deleterious mutations beneficial, because there is a greater density of deleterious than beneficial mutations in the distributions.

Last, we constructed a 95% confidence interval (CI) for each mutation given its mean selection coefficient and the estimated SEM_mut_ described above. We then counted the fraction of mutations whose 95% CI excludes zero. This yields a conservative estimate for our parameter of interest, the total fraction of mutations that are deleterious (or beneficial), as it is designed to indicate whether any particular mutation is deleterious (or beneficial), not to estimate the proportion (which does not depend on unequivocally classifying any one individual mutation as neutral or not). Nonetheless, we report this value as a conservative lower bound on the estimate of non-neutral mutations.

---

**Figure S1.**
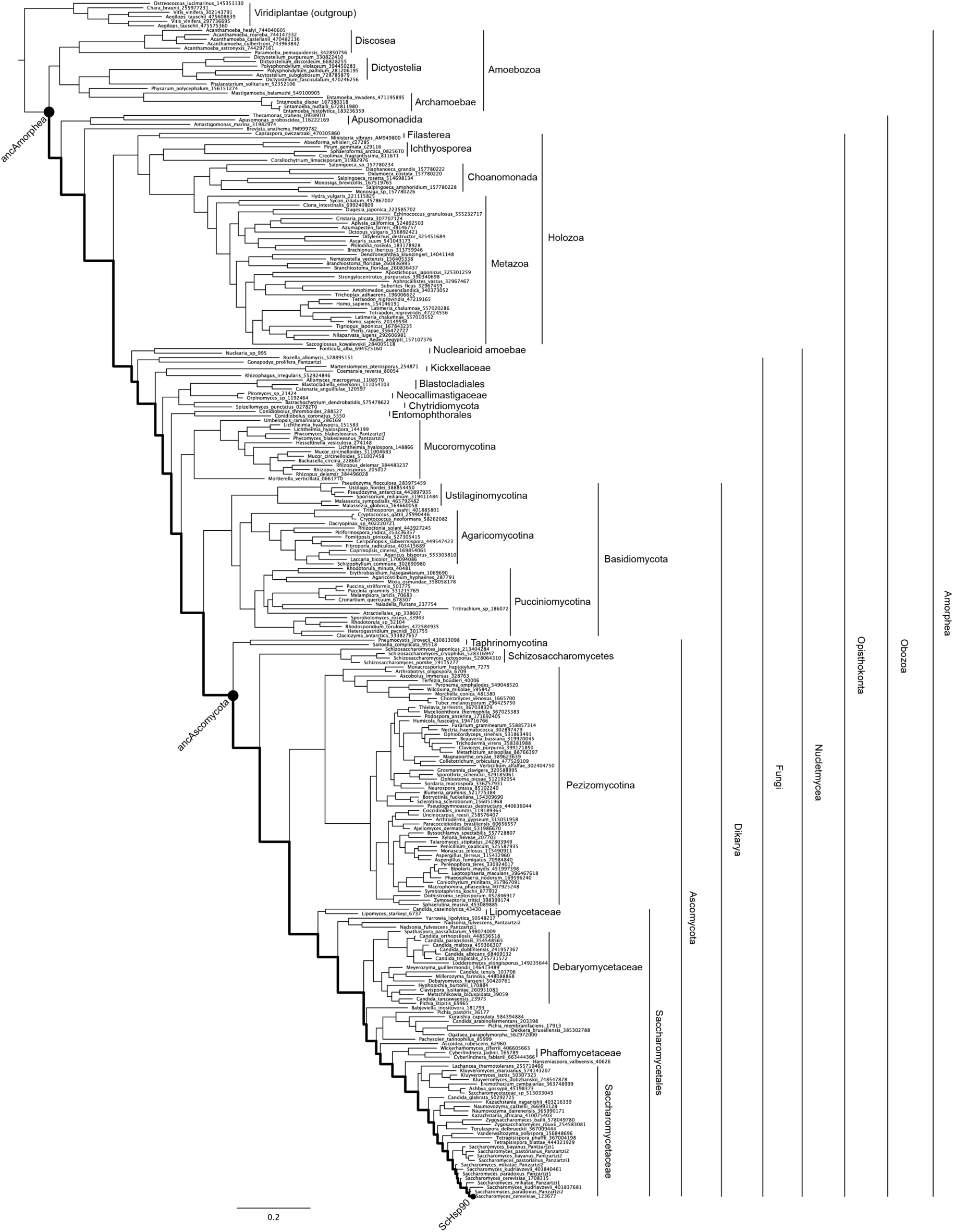
Hsp90 phylogeny. The maximum likelihood phylogeny of 267 Hsp90 protein sequences, with major taxonomic groups labeled. Taxon names indicate genus, species, and an accession number or sequence identifier; complete sequence identification information is given in Dataset S3. Nodes characterized in this study are shown as black dots; the trajectory studied is shown as a thick black line.

**Figure S2.**
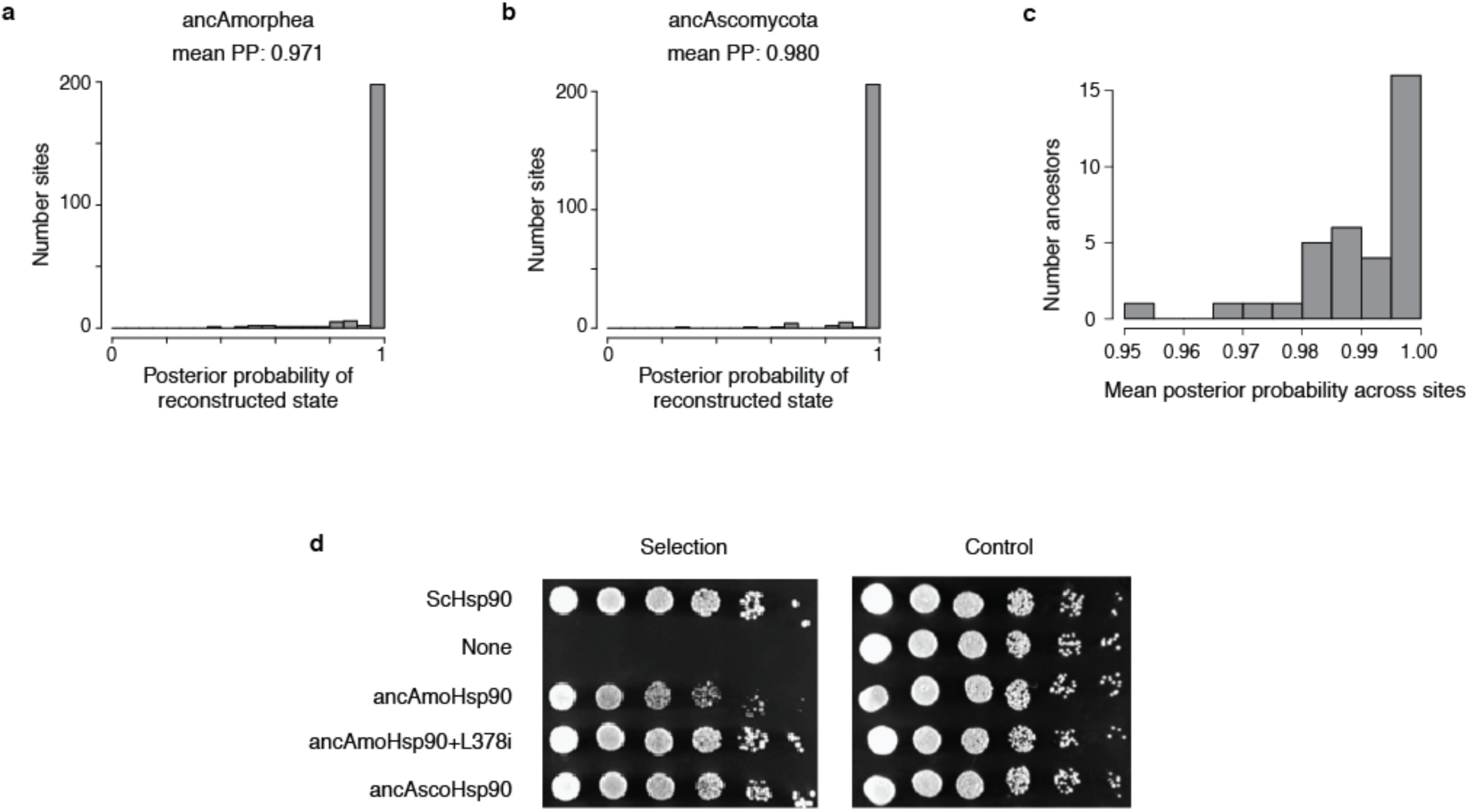
Ancestral Hsp90 sequences have high statistical support and complement yeast growth. **a,b**, For the ancestral NTD sequences reconstructed in this study, the distribution of posterior probability of ancestral states across NTD sites is shown as a histogram. The mean posterior probability of the most probable state across sites (mean PP) is shown for each ancestor. **c**, The distribution of mean PP for reconstructed ancestral sequences along the trajectory from ancAmoHsp90 to ScHsp90. **d**, Growth of *S. cerevisiae* Hsp90 shutoff strains complemented with ancestral Hsp90 NTD variants. Spots from left to right are 5-fold serial dilutions. Control plates represent conditions in which the native ScHsp90 allele is expressed. Under selection conditions, the native ScHsp90 allele is turned off, and growth can only persist when a complementary Hsp90 allele is provided. The ancAmoHsp90 NTD expressed as a chimera with the Sc middle and C-terminal domains exhibits a slight growth defect; this is rescued by adding an additional reversion to the ancAmoHsp90 state in the middle domain (L378i), which occurs on a middle domain loop that extends down and interacts directly with the N-terminal domain and contributes to the NTD ATP-binding pocket. We subsequently refer to ancAmoHsp90+L378i as ancAmoHsp90.

**Figure S3.**
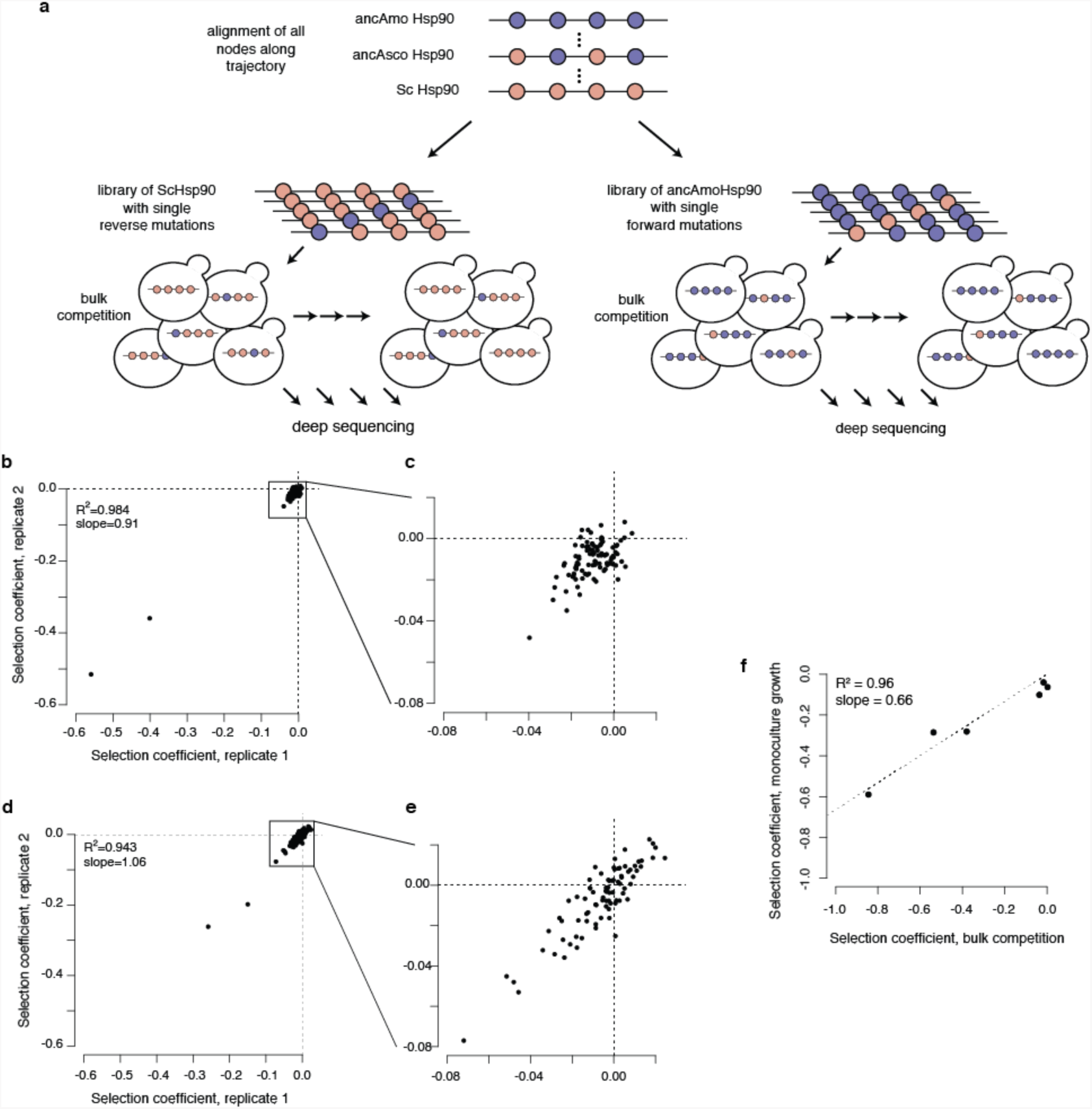
Experimental scheme and reproducibility. **a**, Experimental scheme for testing the fitness effects of individual mutations to ancestral states in ScHsp90 (left) or individual mutations to derived states in ancAmoHsp90 (right). An alignment of all ancestors along the focal trajectory was constructed to identify the trajectory of Hsp90 NTD sequence change from ancAmoHsp90 to ScHsp90. In each background, a library was constructed consisting of the wildtype sequence and all individual mutations to ancestral or derived states. This library was transformed into yeast, which grew through a bulk competition. The frequency of each genotype at each time point was determined by deep sequencing, allowing us to calculate a selection coefficient for each mutation relative to the respective wildtype sequence. **b**, Reproducibility in selection coefficient estimates for replicate bulk competitions of the ScHsp90 library. R^2^, Pearson coefficient of determination. **c**, For visual clarity, zoomed in representation of the boxed region in (**b**). **d**, Reproducibility in selection coefficient estimates for replicate bulk competitions of the ancAmoHsp90 library. R^2^, Pearson coefficient of determination. **e**, For visual clarity, zoomed in representation of the boxed region in (**d**). **f**, Correlation in fitness as measured via bulk competition or monoculture growth assay. R^2^, Pearson coefficient of determination. The line was forced to go through (0, 0); when freely fit, the intercept term was not significantly different from zero.

**Figure S4.**
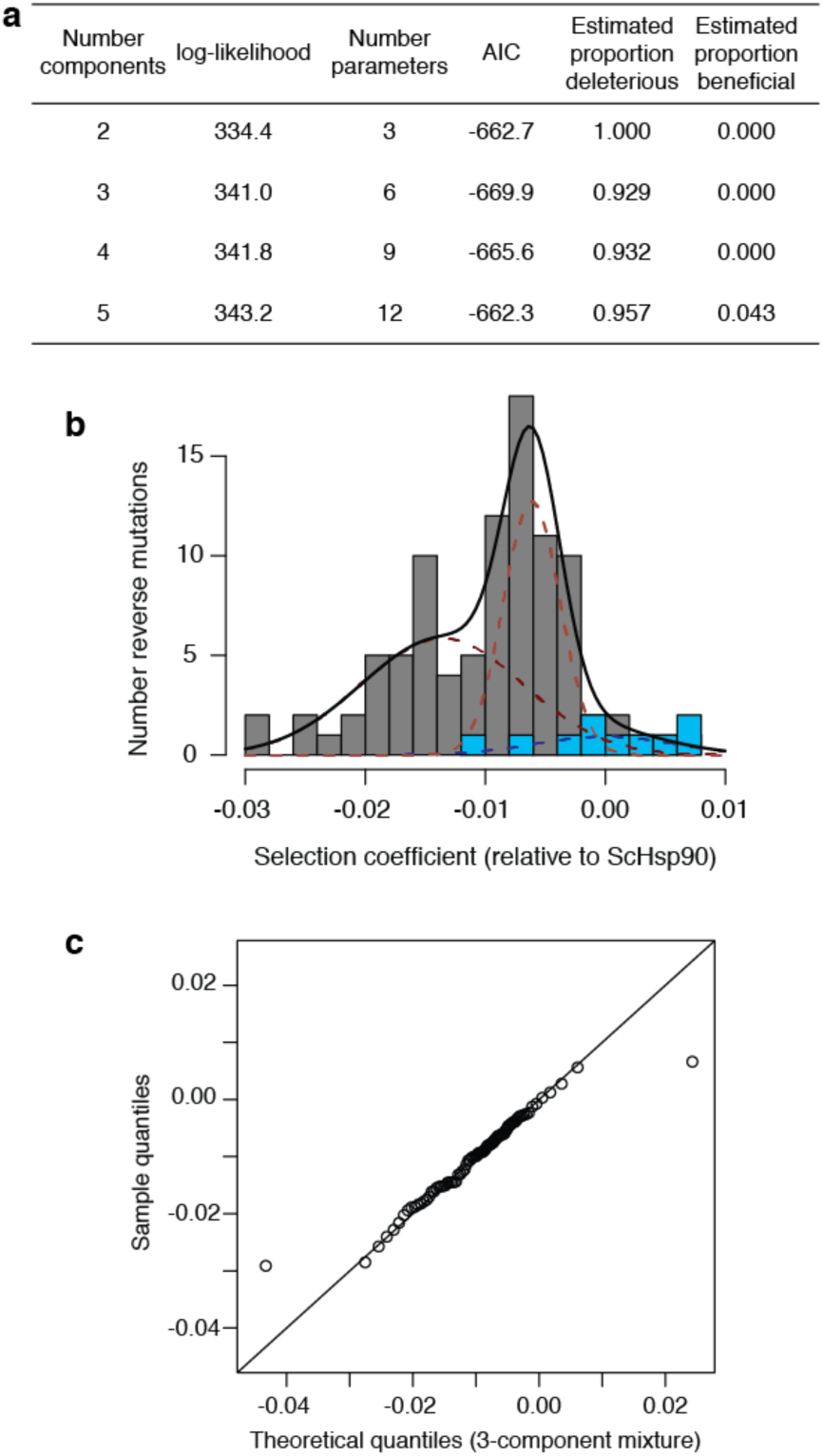
Estimating the proportion of mutations to ancestral states that are deleterious with a mixture model. **a**, Observed selection coefficients of reversions were fit to mixture models containing a variable number of Gaussian distributions; in each case, one distribution is fixed to have the mean and standard deviation of the sampling distribution of independent wildtype ScHsp90 sequences present in the library, the mixture proportion of which is a free parameter; each additional mixture component has a free mean, standard deviation, and mixture proportion. The empirical data were best fit by a 3-component mixture model, as assessed by AIC. Estimated proportion deleterious (beneficial) comes from summing the mixture proportions of components centered below (above) zero. **b**, The best-fit mixture model. Gray bars, observed distribution of selection coefficients of ancestral reversions; blue bars, distribution of observed selection coefficients of wildtype ScHsp90 sequences present in the library. Black line, best-fit mixture model; red dashed lines, individual mixture components centered below zero; blue dashed line, wildtype mixture component. The area under the curve for each mixture component corresponds to the proportion it contributes to the overall mixture model. **c**, Quantile-quantile plot showing the quality of fit of the 3-component mixture model (*x*-axis) to the empirical distribution of selection coefficients of ancestral reversions (*y*-axis). The mixture model assigns more extreme selection coefficients to the tails than is observed in the empirical distribution, but provides a reasonable fit along the bulk of the distribution.

**Figure S5.**
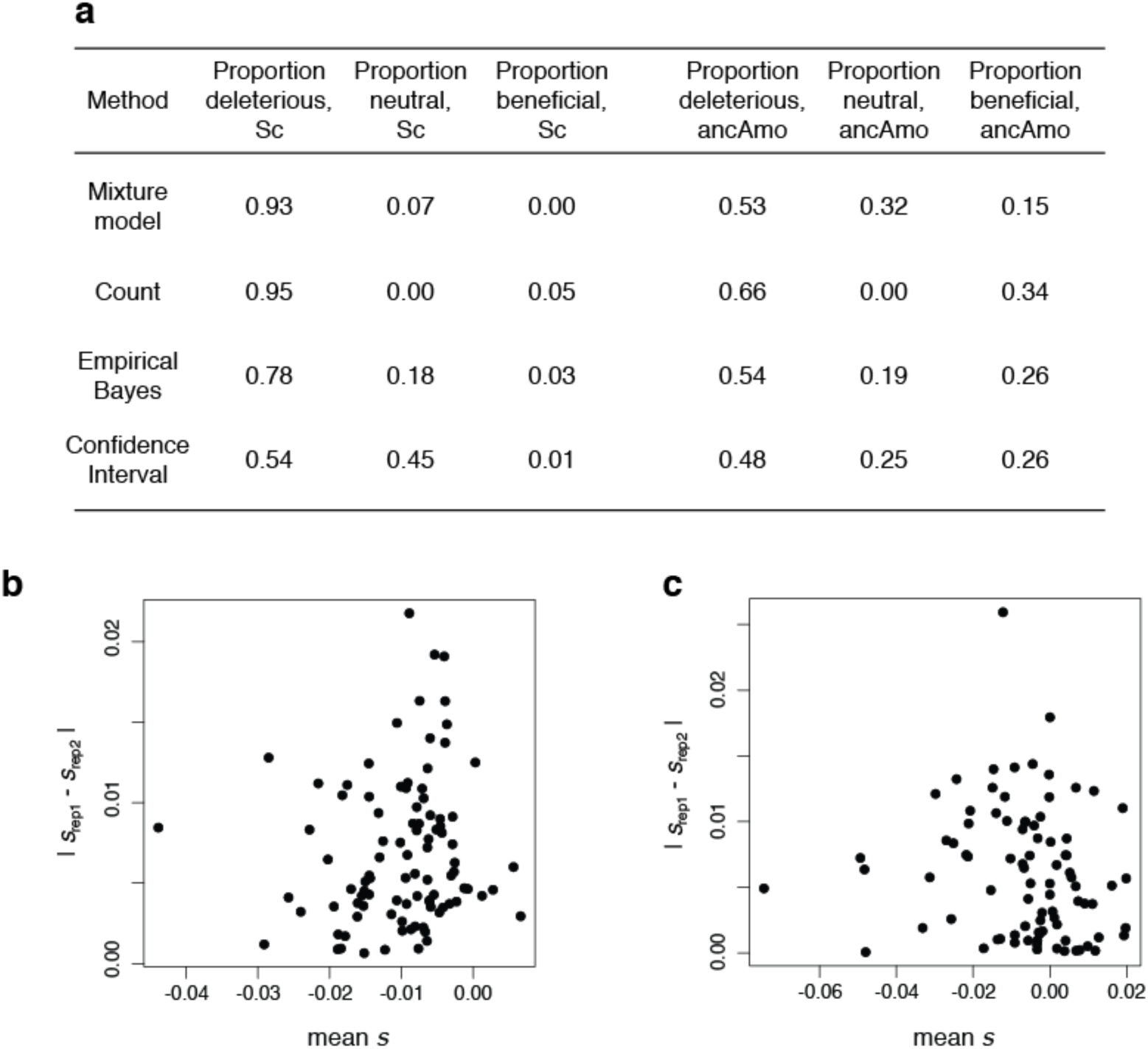
Alternate approaches for estimating the proportion of mutations that are deleterious. **a**, The estimated proportion of mutations that are deleterious, neutral, or beneficial in each background, as determined by each of four statistical methods. See SI Methods for descriptions of each method. **b,c**, Experimental errors are unbiased with respect to the observed selection coefficient. For the ScHsp90 (**b**) and ancAmoHsp90 (**c**) backgrounds, the absolute difference in *s* as determined in each replicate is shown versus their mean. In each background, there is no significant linear relationship between experimental error and *s*_obs_ (*P* = 0.27 and 0.24, respectively, Pearson’s correlation).

**Figure S6.**
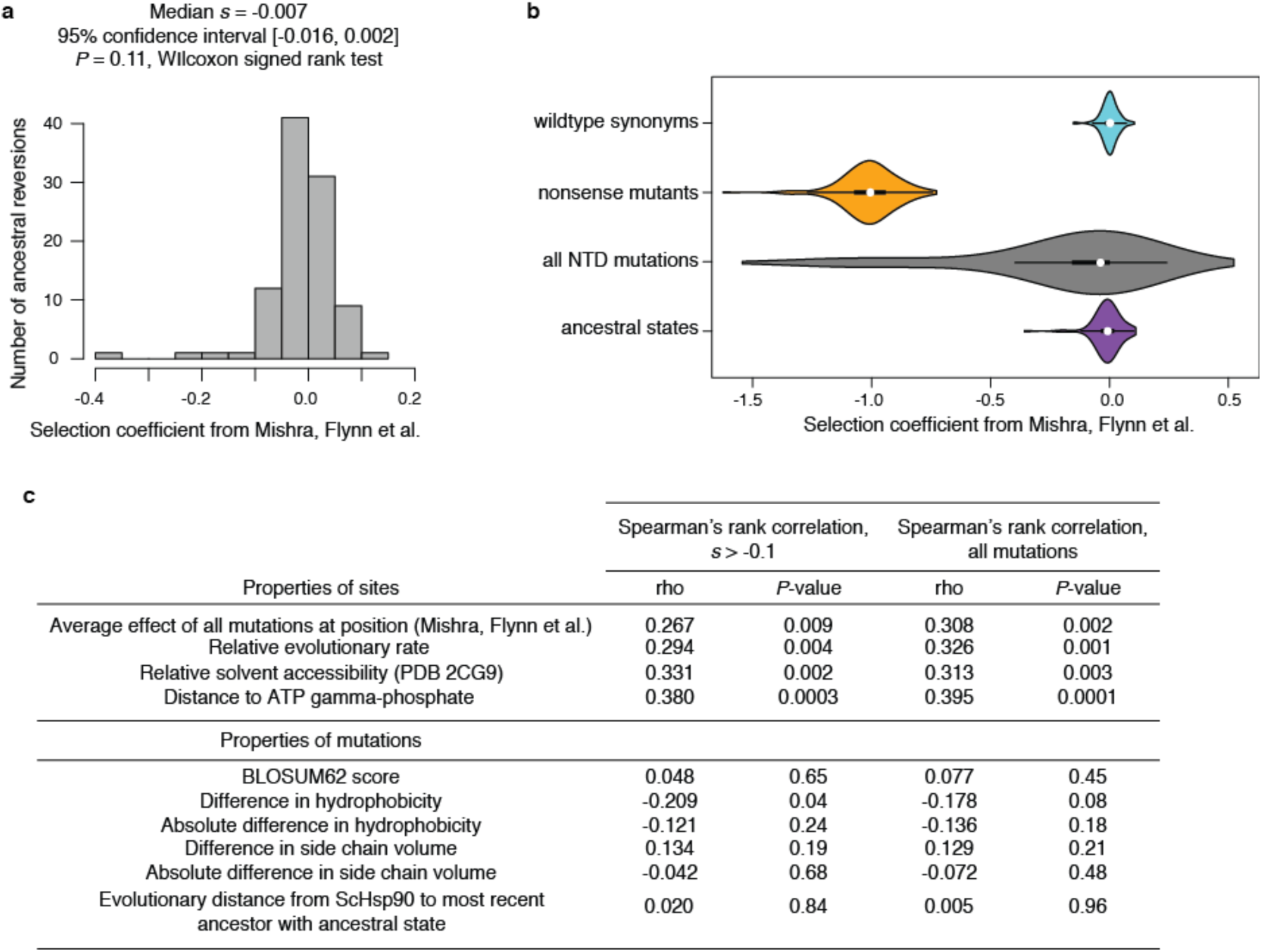
Ancestral states are deleterious in yeast Hsp90. **a**, The signature of deleterious ancestral states is present in the independent but lower-resolution dataset of Mishra, Flynn et al. (46). For each mutation to an ancestral state, the selection coefficient as determined by Mishra, Flynn et al. is shown. The median selection coefficient is −0.007, close to that estimated in the current study; however, this median selection coefficient is not significantly different than zero (*P* = 0.11). Because Mishra, Flynn et al. tested a much larger panel of mutations (all single mutations across the entire NTD), experimental variability of estimated selection coefficients was much larger, possibly explaining the lack of significance of this result in this dataset. **b**, Violin plots show the distribution of mutant effects in the dataset of Mishra, Flynn et al. (46). Ancestral states are less detrimental than the average random mutation in the NTD (*P* = 3.5×10^−9^, Wilcoxon rank sum test with continuity correction). **c**, Reversions exhibit properties typical of genuinely deleterious mutations. For various properties of sites at which we measured the fitness of ancestral variants (top) or properties of the specific amino acids mutated (bottom), we asked whether there was a significant correlation between the property and the selection coefficients of mutations via Spearman’s rank correlation. Ancestral states tend to be more deleterious at positions that are less robust to any mutation, evolve more slowly, are less solvent accessible, and are closer to the gamma-phosphate of bound ATP. These properties are not completely independent; for example, there is a significant positive correlation between relative solvent accessibility and distance to ATP gamma-phosphate. Biochemical properties particular to the amino acid states in each mutation are generally not significantly correlated with the selective effect. Furthermore, we see no evidence for older states being more entrenched, as has been observed by others (1, 4, 14).

**Figure S7.**
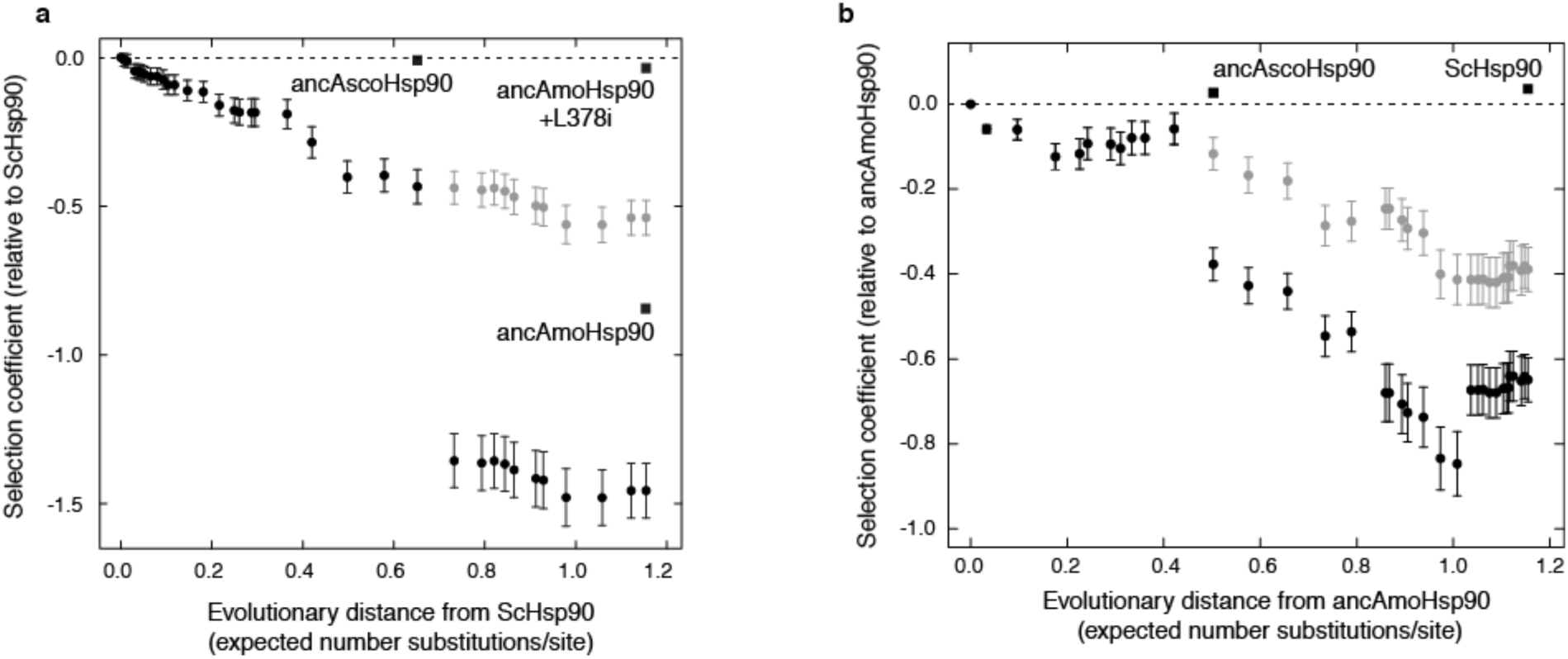
Fitness effects of historical substitutions are modified by intramolecular epistasis. Each black circle represents an ancestral protein along the trajectory from ancAmoHsp90 to ScHsp90. Position along the *x*-axis shows the evolutionary distance that separates it from ScHsp90 (**a**) or ancAmoHsp90 (**b**); *y*-axis position shows the predicted selection coefficient assuming no epistasis relative to ScHsp90 (**a**) or ancAmoHsp90 (**b**). Predicted selection coefficients were calculated as the sum of individual selection coefficients for all sequence differences present in its sequence as measured in ScHsp90 (**a**) or ancAmoHsp90 (**b**). Error bars show the 95% confidence interval for the predicted value, calculated by propagating the standard errors of individual site-specific selection coefficient measurements. Light gray dots show the same data, but excluding the effects of the two strongly deleterious outliers in each library. Labeled squares indicate experimentally determined selection coefficients for complete genotypes: ancAscoHsp90, ancestral Ascomycota (fitness determined via monoculture growth); ancAmoHsp90, ancestral Amorphea (fitness determined via bulk competition); ancAmoHsp90+L378i, ancAmoHsp90 with a candidate epistatic substitution in the Middle Domain also reverted to its ancAmorphea state (fitness determined via bulk competition). Dashed line, *s* = 0.

**Figure S8.**
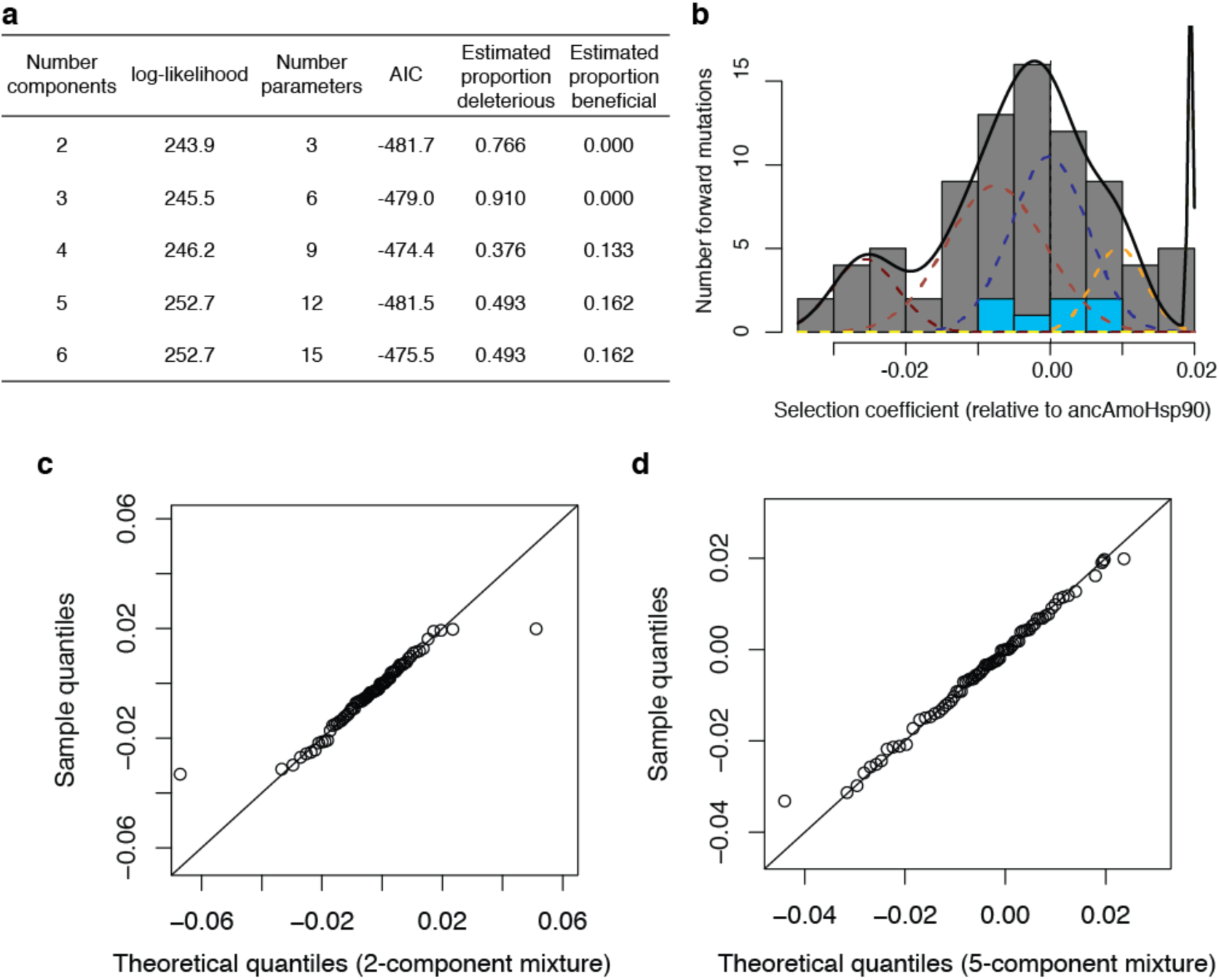
Estimating the proportion of mutations to derived states that are deleterious with a mixture model. **a**, The distribution of selection coefficients of mutations to derived states was fit by mixture models containing a variable number of Gaussian distributions; in each case, one distribution is fixed to have the mean and standard deviation of the sampling distribution of independent wildtype ancAmoHsp90 alleles in the library, the mixture proportion of which is a free parameter; each additional mixture component has a free mean, standard deviation, and mixture proportion. The empirical data were best fit by a 2-component mixture model, as judged by AIC, with a 5-component mixture being almost equally well fit; the 5-component mixture resulted in a more conservative estimate of the proportion of mutations that were deleterious than the 2-component mixture, and so was chosen despite the AIC difference of 0.2. Estimated proportion deleterious (beneficial) comes from summing the mixture proportions of components centered below (above) zero. **b**, The fit of the 5-component mixture model. Gray bars, distribution of selection coefficients of mutations to derived states; blue bars, distribution of selection coefficients of independent ancAmoHsp90 alleles present in the library. Black line, five-component mixture model. Red dashed lines, individual mixture components centered below zero; blue dashed line, wildtype mixture component; yellow dashed lines, individual mixture components centered above zero; relative integrated areas of mixture components correspond to the relative proportions they contribute to the overall mixture model. **c,d**, Quantile-quantile plot showing the quality of fit of the 2-component (**c**), or 5-component (**d**), mixture models (*x*-axis) to the empirical distribution of selection coefficients of mutations to derived states (*y*-axis).

**Figure S9.**
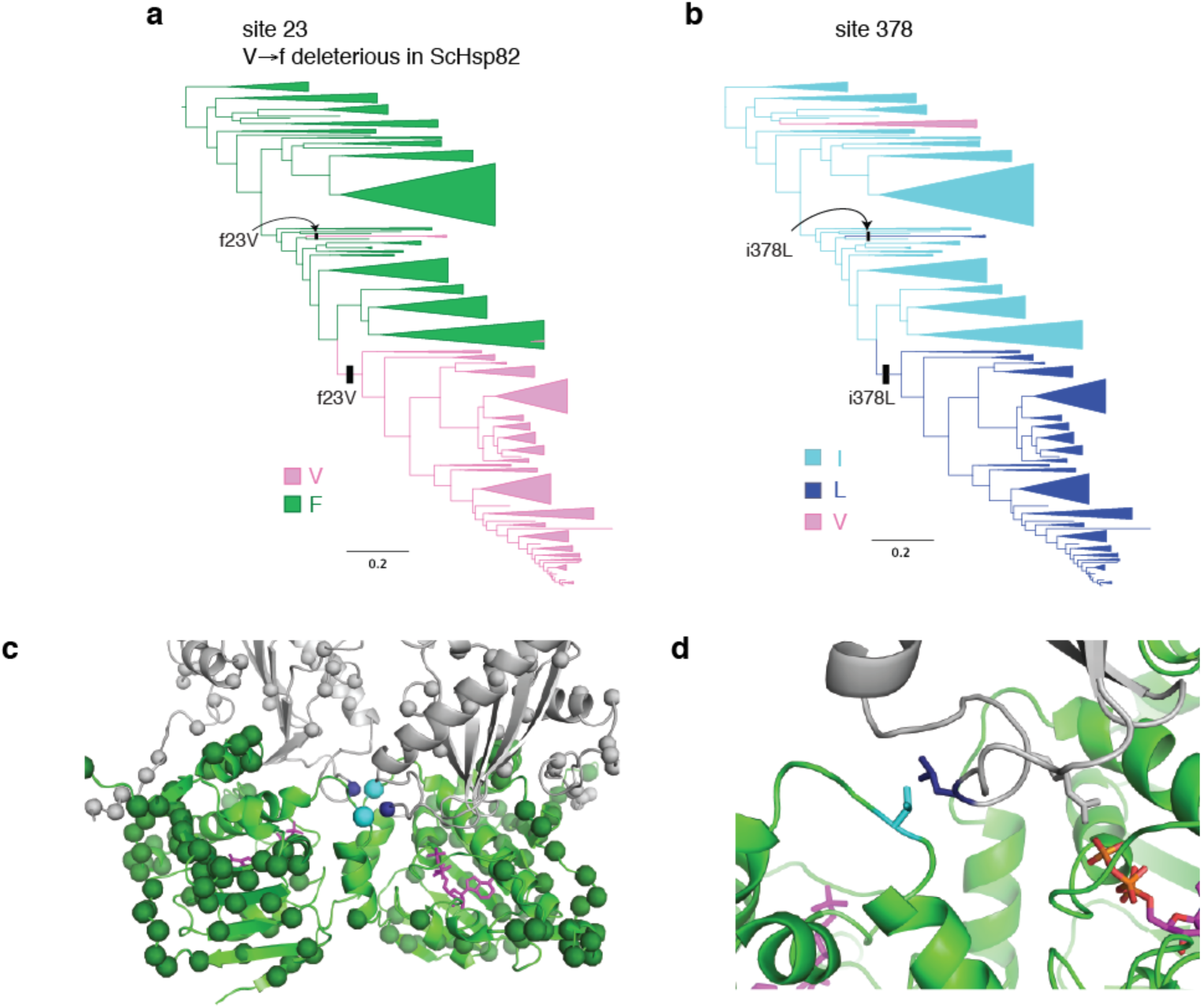
The deleterious V23f reversion is ameliorated by L378i. **a,b** Character state patterns at sites 23 (**a**) and 378 (**b**). On the lineage to ScHsp90, f23V co-occurred with i378L before the common ancestor of Ascomycota. The same two substitutions also co-occur on an independent lineage on this phylogeny (Kickxellaceae fungi), and in the distantly related Rhodophyta red algae (not shown). **c**, The locations of sites 23 and 378 on the ATP-bound Hsp90 dimer structure (PDB 2CG9). Cyan spheres, site 23; dark blue; site 378; dark green, other variable NTD sites; gray, other variable middle and C-terminal domain sites. Magenta sticks, ATP. **d**, Zoomed view of sites 23 and 378. These side chains are in direct structural contact, and may be important for the positioning of the middle domain loop that bears R380 (gray sticks), which forms a salt bridge with the ATP gamma-phosphate and is critical for ATP binding and hydrolysis (34, 58).

**Figure S10.**
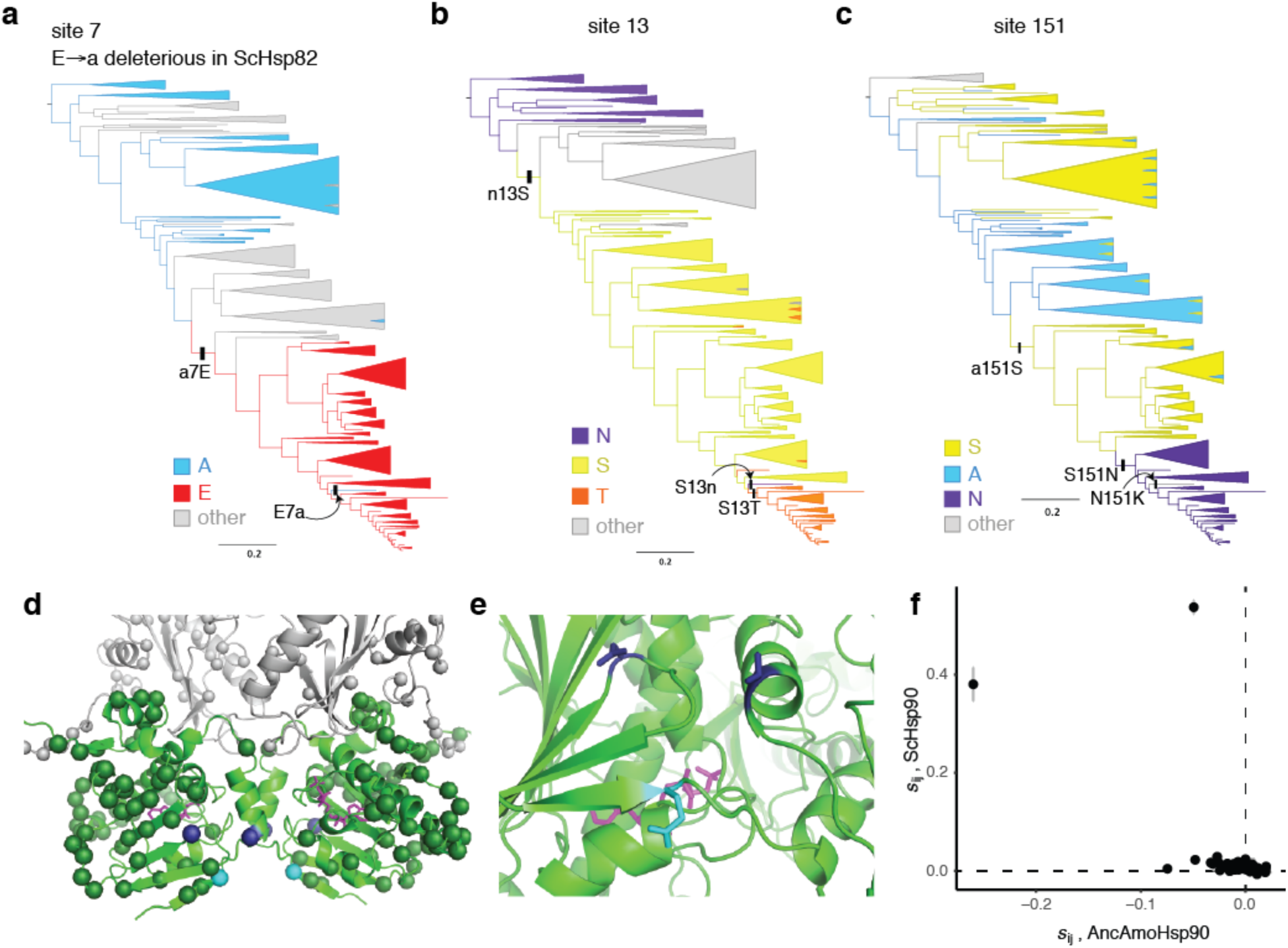
The deleterious E7a reversion is partially ameliorated by N151a or T13n. **a,b,c**, Character state patterns at sites 7 (**a**), 13 (**b**) and 151 (**c**). On the trajectory to ScHsp90, a7E occurred before the common ancestor of Ascomycota, then later reverted in the lineage leading to *Ascoidea rubescens* (arrow); on this latter lineage, site 13 also reverted to the ancestral state asparagine, and site 151 substituted to a third state lysine. **d**, The locations of sites 7, 13, and 151 on the ATP-bound Hsp90 structure (2CG9), represented as in Fig. S9c. Cyan spheres, site 7; dark blue, sites 13 and 151. **e**, Zoomed in view of sites 7, 13, and 151. These side chains are not in direct physical contact; however, site 7 is on a beta strand that undergoes extensive conformational movement when Hsp90 converts between ADP-and ATP-bound states. **f**, The same plot as Fig. 5c is shown, including the two strongly outliers V23f and E7a. See Fig. 5c legend for details.

